# Phylogeny, thermal seasonality, and life history shape trait variation in annual and perennial barley relatives

**DOI:** 10.64898/2026.06.02.729615

**Authors:** Michael Anokye, Timo Hellwig, Einar Baldvin Haraldsson, Rebekka Schüller, Nina Döring, Philipp Westhoff, Anna Bucharova, Maria von Korff

## Abstract

- There is growing interest in developing perennial cereal crops for sustainable production, yet key differences in trait syndromes between annual and perennial grasses and their influence on environmental adaptation remain poorly understood.
- We measured 25 traits in 16 annual and perennial *Hordeum* species (45 accessions), including barley, grown over three seasons in a common garden. Using a phylogenetic framework and repeated transitions between annual and perennial forms, we (i) identified traits distinguishing these life strategies and (ii) tested how they relate to climate at the accessions’ origins.
- Wild and cultivated barley are distinguished within the *Hordeum* clade by high growth rates and large organs, which may have predisposed wild barley to domestication. Annual and perennial accessions differed in resource allocation: annuals had higher harvest index and leaf and grain nitrogen, while perennials produced carbon-rich tissues and sustained vegetative growth. Seasonal temperature variation shaped trait syndromes: annual traits aligned with temperature in the driest quarter, reflecting selection under terminal stress, while perennial traits correlated with temperature in the wettest quarter, the main growth phase shaping long-term performance and survival.
- We provide important information on traits and climate adaptations underlying perennial persistence and annual resource strategies, relevant for developing perennial cereal crops.

## Introduction

Variation in life history represents a key adaptive strategy in plants to optimize survival and reproduction in different environments (Stearns, 1992; Sultan, 2000; Friedman, 2020). Within many taxa, the allocation and distribution of resources to reproduction within seasons versus between seasons is very variable, with life spans ranging from a few weeks in short-lived herbs to hundreds of years in long-lived trees (Friedman, 2020). While the diversity in plant life histories has made classification challenging, the most widely used classifications are based on seasonal life expectancy, e.g., annual, biennial, and perennial (Friedman, 2020), or the number of reproductive bouts during life, monocarpy/semelparity, polycarpy/iteroparity (Kirkendall & Stenseth, 1985; Silvertown, 1989; Hughes & Simons, 2014; Hughes, 2017). Annual monocarpic species complete germination, fertilization, and grain release within one season, then undergo whole plant senescence and survive the unfavorable season in the form of grains (Zanne et al., 2014; Boyko et al., 2023). On the contrary, perennial species survive and reproduce over multiple seasons, and vary in their life expectancy, from biennial plants that die after one year of vegetative and a second year of reproductive growth (monocarpic) to perennials that can survive many years, reproducing each year (polycarpic; Friedman, 2020; Zhao & Wang, 2024). The advantages of either strategy depend on the relative benefits of immediate reproduction balanced against survival and future reproduction. Annuals are favored when adult plant survival is low, and grain persistence and seedling survival are high. Consequently, annuals prevail in unpredictable, strongly fluctuating environments with prolonged periods of harsh conditions, including seasonally high temperatures and drought, compared to perennials (Stearns, 1992; Poppenwimer et al., 2023).

Annuality versus perenniality is a trait syndrome that combines traits related to resource use, growth, and reproductive timing. Annual plants optimize their reproductive effort through high resource acquisition and rapid growth, whereas perennial plants have evolved strategies that emphasize long-term survival and resource conservation (Grime & Hunt, 1975; De Souza et al., 1987; Garnier, 1992; Lindberg et al., 2020). Most studies have suggested that annuals exhibit higher relative growth rates and biomass production, and a larger allocation of resources to reproductive structures and specific root length (Grime & Hunt, 1975; De Souza et al., 1987; Garnier, 1992; Roumet et al., 2006). Perennials, by contrast, are characterized by traits that allow for survival and stress resistance, such as high above-ground and below-ground tissue density and allocation of a higher proportion of biomass into roots (Roumet et al., 2006; Gonzalez-Paleo et al., 2024). As such, annual and perennial species differ in major functional traits and trait complexes that represent compromises between diverging selection pressures resulting from combinations of competition, stress, and disturbance (Grime, 1977).

Life history variation within the Pooideae (temperate grasses), which includes major crop species such as wheat (*Triticum* L. ssp.) and barley (*Hordeum vulgare* L.), is prevalent, with many independent transitions between annual and perennial life forms in different lineages (Lindberg et al., 2020). Variation in growth performance is closely linked to the leaf economics spectrum, which describes coordinated relationships among leaf traits—including specific leaf area (SLA) and leaf nitrogen (N) and carbon (C) contents, thereby representing a gradient from conservative to acquisitive plant strategies (Wright et al., 2004; Joswig et al., 2022). Typically, annual species exhibit rapid growth and build leaves that enable greater carbon capture per unit leaf area (higher SLA), but with lower leaf tissue density and higher N content. In contrast, sister perennial species are characterized by higher leaf density and thickness (i.e., lower SLA) with relatively low N content, which is important for longevity and persistence (Garnier, 1992; Garnier & Laurent, 1994; Garnier et al., 1997). These consistent differences in leaf-specific traits between annuals and perennials were interpreted as adaptive for the two life-history strategies and linked to fast growth and high productivity in annuals and slow growth and persistence in perennials (Garnier, 1992; Garnier & Laurent, 1994). Furthermore, Lindberg et al. (2020) recently suggested that high relative growth rates and the preferential allocation of resources to above-ground structures represent evolutionary precursors of transitions from perennial to annual life history strategies. However, several studies on congeneric annual and perennial pairs of grass species have only confirmed a consistent difference in growth and resource allocation during early development, but not in older plants (Garnier, 1992; Atkinson et al., 2016). Similarly, studies contrasting domesticated annual and perennial wheat and rye species found limited support for the hypothesis that annuals have higher carbon assimilation rates (Jaikumar et al., 2014). These findings suggest that the physiological and morphological differences between annual and perennial species are complex and may be confounded by evolutionary and ecological factors that affect trait variation.

The genus *Hordeum* belongs to the economically important grass tribe Triticeae and comprises 33 species, including diploids and polyploids (Brassac & Blattner, 2015). The species are disjunctly distributed in arid and temperate areas of the Northern Hemisphere, South America, and South Africa (Blattner, 2006; Blattner et al., 2010; Bothmer et al., 1995). *Hordeum* also includes the annual crop barley (*H. vulgare* subsp. *vulgare*), one of the earliest domesticated cereal crops selected from its wild progenitor *Hordeum vulgare* subsp. *spontaneum* approximately 10,000 years ago in the Fertile Crescent (Zohary & Hopf, 1993; Badr et al., 2000). Domestication and improvement breeding of barley were accompanied by a suite of morphological and life history changes that enhanced harvestability and yield (Haas et al., 2019). Wild barley has been widely used to introduce beneficial traits and to study the genetics of domestication, but apart from *H. bulbosum*, other wild relatives remain largely unexplored as potential breeding resources due to limited genetic and genomic tools and knowledge of their morphological and physiological diversity. The recent establishment of reference genomes for 27 *Hordeum* species and additional characterization of intraspecific variation now allows us to disentangle convergent evolution of life history traits from phylogenetic dependency in *Hordeum* (Hellwig et al., 2025; Feng et al., 2025).

The growing interest in developing perennial cereal crops for sustainable food production has prompted research to dissect the morphological, physiological, and genetic differences between annual and perennial grasses (Crews et al., 2018). While the physiological and genetic mechanisms controlling development in annual model and crop species have been extensively studied over the past decades, a limited understanding of the mechanisms underlying differences in life history strategies has hindered progress in perennialising annual crops.

Here, we build on an improved phylogeny to systematically study 25 plant functional traits in 16 *Hordeum* species (45 accessions) in a common garden. Our objective was to identify key traits and trait combinations that distinguish annual and perennial *Hordeum* species and to explore the extent to which parallel trait syndromes have been deployed during the repeated transitions between perennial and annual *Hordeum* species. We explored whether life history differences in *Hordeum* follow the acquisitive–conservative axis of plant ecological strategies. We hypothesized that the long-lived perennials are slow-growing with late investment in reproduction, while annual *Hordeum* accessions grow faster, have earlier and greater reproductive effort. In this context, we tested whether the leaf economic spectrum, measured as leaf trait covariation, captures the balance between leaf construction costs and plant growth potential and is linked to variation in resource allocation patterns, growth, and longevity, thereby varying between annual and perennial species. Finally, we explored whether climatic variables at the accessions’ origins are linked to trait variation differentiating annual from perennial accessions across the *Hordeum* phylogeny.

## Materials and Methods

### Plant material

A total of 45 accessions, belonging to 16 *Hordeum* species, with between one and six accessions per species, were used in the study (Table S1, Fig. 1a,b). These included three diverse wild barley accessions (*Hordeum vulgare* ssp. *spontaneum*) from Israel, Turkey, and Iraq (Badr et al., 2000; Liller et al., 2017) and three barley cultivars (*Hordeum vulgare* ssp. *vulgare*). The barley cultivars included a two-rowed winter barley cultivar (cv Igri), a two-rowed spring cultivar (cv Golden Promise), and a six-rowed spring cultivar (cv Morex). The remaining wild *Hordeum* accessions originated from Western Eurasia, the Mediterranean, and North and South America (Bothmer et al., 1995). Apart from the barley cultivars, all accessions were obtained from genebanks and are documented as diploid species with either an annual or perennial life history, except *Hordeum murinum* subsp. *murinum*, which is a tetraploid.

**Fig. 1:**
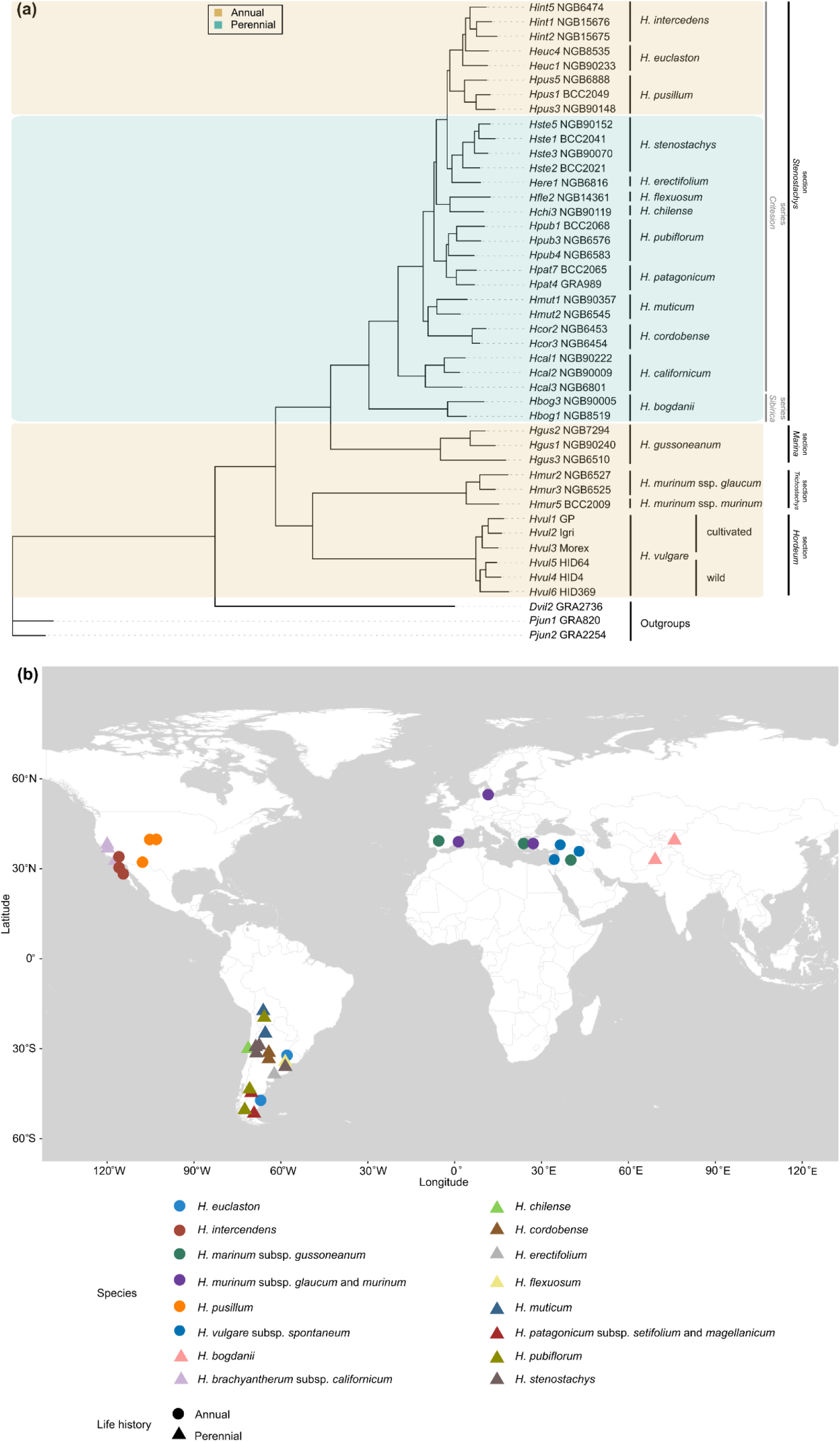
Accession-based phylogenetic tree and map of the accessions’ origin. **(a)** Accession-based phylogenetic tree showing 41 accessions included in the linear model (excluding outgroups). Rectangles drawn around accessions/species are colored by life history: brown for annuals (n = 20) and teal for perennials (n = 21). Infrageneric groupings (section and series) follow the classification of Blattner (2009). Tree modified from Hellwig et al. (2025). **(b)** The map shows the geographic distribution of accessions for each species. Colors on the map represent different accessions within species, and shapes indicate life history groups (annuals, n = 17; perennials, n = 21). Barley cultivars (Morex, Igri, and Golden Promise) were excluded from the plot.

### Growth conditions and experimental design

For all *Hordeum* accessions, grains were pre-germinated in 96-well trays manually filled with a freshly prepared soil mixture consisting of 99.6% (v/v) Mini tray soil (MIM800, Balster Einheitserdewerk, Fröndenberg, Germany) and 0.4% (v/v) Osmocote Exact Standard 3-4M (Scotts Company LLC). Trays were then placed in a controlled-environment growth chamber, where grains were stratified for 14 days at 4 °C in the dark to promote uniform and synchronous germination. After stratification, the trays were transferred to long-day (LD) conditions (16h light/8h dark, 22 °C/16 °C) in a controlled-environment growth chamber. Seedling emergence was scored and ranged from 5 to 10 days after transfer to LD growing conditions. Once seedlings had developed two to three leaves, they were moved to short-day (SD) conditions (8h light/16h dark) and 4 °C (day and night) and vernalized for 12-14 weeks before transplanting to the experimental area at the Botanical Garden (BG) of Heinrich Heine University (HHU), Düsseldorf, Germany (51°11’10.4“N 6°48’09.9”E). After vernalization, individual seedlings were repotted into 7.5 L pots (25.9 cm x 20.6 cm), manually filled with 7 L of soil mixture containing 93 % (v/v) Einheitserde ʻED73’ peat soil (Einheitserde Werkverband e.V., Sinntal Altengronau, Germany), 6.6 % (v/v) sand, and 0.4 % (v/v) Osmocote exact standard 3-4M (Scotts Company LLC). The seedlings were arranged in a randomized complete block design with 4 blocks and 4 plants per accession in each block, in total 16 plants per accession. During the growth period, water was manually provided 2-3 times a week, except on rainy days. No additional fertilization, disease, or pest management practices were applied. The air temperature at the experiment area was monitored on-site (Table S2) using a Spectrum WatchDog 2900ET Weather Station (Spectrum Technologies, Inc., Fairborn, USA). The experiment was repeated in 2021, 2022, and 2023, from March/April to the end of August/September.

## Phenotyping

### Shoot and grain phenotyping

In total, 25 continuous vegetative and reproductive traits were measured in four biological replicates per accession across three experimental seasons (Table S3). After 30 days of plant establishment in the field, the growth rate was assessed by harvesting and measuring the total aboveground dry shoot biomass. Days to flowering were determined as the days from transplanting to the outdoor pots until the first awns from the flag leaf appeared, Zadok stage 49 (Zadoks et al., 1974). The remaining traits were evaluated at the grain-filling stage and included flag leaf length, width, and area; plant height; floret number; and spike length. Unless stated otherwise, for all traits measured at grain filling, the main tiller was tagged per plant. The length of the flag leaf blade was measured from the leaf collar to the blade’s tip. The width of the same flag leaf was measured at the widest section of the blade. The length of the main stem spike was measured from the collar at the junction of the stem and the spike to the spike tip, excluding the awns. Plant height was measured from the base of the main tiller at the soil surface to the collar of the spike. The floret number per spike was manually counted along the main tiller spike. One-sided area of an individual leaf (leaves rehydrated for 24 hrs in 50 ml tubes) of the main tiller flag leaf was determined using the Petiole application (non-pro version 4.0.2; Singh et al., 2021; Kizildeniz, 2023) after calibration (https://petioleapp.com/pads/Petiole-calibration_pad_5.pdf). Flag leaf area and dry weight were used to calculate specific leaf area (SLA) by dividing the flag leaf area (cm^2^) by the dry weight (mg) after drying at 65 °C for 72 h.

At harvest, the stem diameter at the base of the main tiller was measured on the mid-sections between the first and second nodes above the soil surface using a digital caliper (Futec DC-1, Futec, Austria; resolution 0.01 mm; accuracy ±0.02 mm). Further, total dry shoot and reproductive biomass represented by spike weight were quantified post-harvest. Harvest index was calculated as dry spike biomass divided by total dry shoot biomass at harvest. Grain traits were measured on all grains from 3–4 mature spikes per plant (*c.* 30–50 grains) using a MARVIN Seed Analyzer (GTA Sensorik GmbH, Germany), which quantified grain length, width, length-to-width ratio, area, circularity, and thousand-grain weight (TGW). We also scored regrowth and winter survival to confirm the annual and perennial growth habits as described by Bothmer et al. (1995). All annual accessions underwent whole-plant senescence at the end of the season. In addition to the repeated sowing over three years, we tested for regrowth and flowering for at least 2 seasons in at least 1 accession of each perennial species (Table S1). For the remaining perennial accessions, survival in subsequent years was not recorded because of seed contamination between nearby pots. Thus, we confirmed the classification of annual and perennial species by Bothmer et al. (1995) in our common-garden conditions.

### Measurement of leaf and grain carbon and nitrogen content

We used flag leaf tissues collected from the three oldest reproductive tillers at the grain-filling stage, and corresponding grain samples from four biological replicates per accession for carbon (C) and nitrogen (N) measurements. Flag leaf material was shredded into a 2 ml reaction tube containing two (3 mm) steel beads, and dried at 65 °C for 72 h in a drying and heating cabinet. Dried leaves were then ground to a fine powder using a TissueLyser II (QIAGEN GmbH, Hilden, Germany).

Grains (c. 50-100 grains of wild *Hordeum*, c. 30-50 grains of wild and cultivated *H. vulgare*) were manually pulverized using a ceramic mortar and pestle, and 0.04 g of the pulverized sample was transferred to a 2 ml reaction tube with two (3 mm) steel beads, dried overnight at 65 °C, and ground to a fine powder. C and N concentrations in 2 mg of homogenized leaf and grain powder were determined using the Isoprime 100 isotope ratio mass spectrometer coupled to an isotope cube elemental analyzer (Elementar, Hanau, Germany), adapting standard procedures previously described (Coplen et al., 2006; Gowik et al., 2011). All C and N measurements were done at the Center of Excellence on Plant Sciences, Metabolomics and Metabolism Laboratory (CMML, HHU, Düsseldorf, Germany).

### Phylogeny

Phylogenetic relationships among accessions were derived from the accession-based phylogenetic tree published by Hellwig et al. (2025). This tree was pruned to retain only the accessions included in the present study. Four accessions, *H. euclaston* (NGB90342), *H. bogdanii* (NGB90046), and *H. chilense* (GRA972, NGB90118) were not included in the phylogenetic tree, but these species remained represented by other accessions (Fig. 1a, Table S1) and thus excluded in all analyses accounting for phylogenetic structure. Patristic distances among all branch tips were calculated using the ape package (v5.3; Paradis & Schliep, 2019). The resulting phylogenetic distance matrix was subjected to principal coordinate analysis (PCoA; Desdevises et al., 2003), and the first three axes (PCoA1–PCoA3), which together explained approximately 95% of the phylogenetic variation, were retained. These were included as covariates in all subsequent analyses to account for phylogenetic structure.

## Statistical analyses

All trait data were analysed using R (R v.4.4.0; Core Team, 2024). Best Linear Unbiased Prediction (BLUP; Henderson, 1975) across 25 traits and the three seasons (Table S4) were estimated from a fitted mixed linear model (LM) using restricted maximum likelihood with the *lmer* function in lme4 (v.1.1-7; Bates et al., 2015) for each continuous trait:

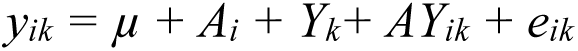

where the response variable *y_ik_* is a function of the overall mean (*µ*), random effects of accession (*A_i_*), year (*Y_k_*), accession-year term (*AY_ik_*), and the residual error term (*e_ik)_*. BLUP-adjusted means for the accessions (A) were extracted using the *ranef* function in lme4 (v.1.1-7; Bates et al., 2015).

PCA was performed on a scaled and centered BLUP-adjusted means of accessions across years for 25 traits (Table S4) using the *prcomp* function in base R. Eigenvalues were extracted to observe the percent variance explained by respective principal components using the *get_eigenvalue* function from the R package factoextra (v.1.0.7; Kassambara & Mundt, 2020). Plots of PCA variables were obtained using the R package ggbiplot (v.0.55; Vu, 2011). PCA loading for the first three principal components (PC1-PC3) was extracted and visualized as a heatmap using the R package pheatmap (v.1.0.13; Kolde, 2025; Fig. S1). Numerical loading values were overlaid on the heat map to summarize the strength and direction of associations. To account for the nonindependence of related species in our ordination, we also ran a PCA on BLUP-adjusted means after phylogenetic correction using phylogenetic generalized least squares (PGLS), implemented via the *pgls* function in the caper package in R (v.1.0.4; Orme et al., 2025). For each trait, our PGLS model assumed a Brownian motion model of trait evolution (Felsenstein, 1985), with Pagel’s λ estimated by maximum likelihood (Pagel, 1999; Freckleton et al., 2002). For both analysis (phylogeny corrected and uncorrected PCA), differences in principal components (PC1 to PC3) between cultivated, wild barley and all other annual and perennial *Hordeum* species were tested via one-way analysis of variance (ANOVA) comparison of median testing using the *aov* based R function (*p* ≤ 0.05) followed by posthoc tests using the *LSD.test* function implemented in the agricolae package in R (v.1.2-3; Mendiburu & Simon, 2015).

To test for significant effects of life history on trait expression, we fitted a set of linear models (LMs) that accounted for phylogenetic relationships using three proxy PCoA axes derived from the phylogenetic distance matrix as covariates. Each trait matrix was regressed against the phylogenetic principal components, and the model residuals were retained as phylogenetically corrected trait values. Effect sizes were calculated for contrasts between individual traits of annual and perennial accessions, using the *cohen_d* function from the effectsize package in R (v.0.8.8; Ben-Shachar et al., 2020). The approach is based on Cohen’s (1988) standardized mean difference framework and Hedges (1981) small-sample bias correction. Confidence intervals (95%) were obtained via percentile bootstrap resampling (1,000 iterations) assuming unequal variances.

To assess differences in trait syndromes between annual and perennial accessions, pairwise trait correlations (with Bonferroni-adjusted *p*-values) were calculated for all accessions and separately for each life-history group using Pearsońs correlation tests implemented in the *cor.test* function from the stats package in R. Results were visualized using the *corrplot* function in the R package corrplot (v.0.84; Wei & Simko, 2017).

### Redundancy analysis (RDA)

To test the effects of climate at the accessions’ origins on trait variation, we conducted redundancy analysis (RDA), a multivariate method for analysing linear relationships between two or more matrices (Legendre & Legendre, 2012). For the RDA analysis, we included all accessions represented in the phylogenetic tree (Fig. 1a,b), except for the barley cultivars Morex, Igri, and Golden Promise, for which no georeferenced climate data were available (Table S1). The RDA analysis was thus conducted on 38 accessions from 16 species. Specifically, we used simple RDA (when species or accessions’ phylogenetic relatedness was not accounted for) and partial RDA conditioned on phylogeny to estimate the proportion of trait variation explained by climatic variables. RDA was performed with the *rda* function of the R package vegan (v.2.5-6; Oksanen et al., 2019). For all RDA models in this study, we performed 5000 permutations to test the significance of explanatory variables using the R function *anova.cca*.

Climate variables were derived from a dataset published by Hellwig et al. (2025), which comprises 13,439 publicly available occurrence records of all 21 diploid *Hordeum* species. For each, 19 bioclimatic variables (Table S5) from WorldClim2, averaged across the years 1970-2000 at a 2.5-minute spatial resolution, were downloaded (Fick & Hijmans, 2017). To account for collinearity among these variables, we identified six mutually uncorrelated groups of highly correlated variables following an approach described by Chang et al. (2022). To identify representative environmental predictors from these, we built climatic niche models using generalized linear models (GLMs) implemented in ENMTools (v.1.0; Warren et al., 2021). Each model included one variable per group, ensuring that no correlated predictors were used together. This resulted in 1,152 candidate models, representing all possible combinations of variables across groups. Model performance was evaluated using Akaike’s Information Criterion (AIC), and the best-fitting model was selected accordingly. The six variables from the best-fitting model (Table S5) were used for downstream analyses because they best captured the occurrence patterns of diploid *Hordeum* species.

We then downloaded data on the six selected bioclimate variables for the 38 analysed accessions herein, based on their geographic origins, averaged across the years 1970-2000 at 2.5-minute spatial resolution (Fick & Hijmans, 2017). We used a combination of all six climate variables, and single climate variables, in simple and partial RDA models. To evaluate the effects of individual climatic variables on trait variation, we sequentially fitted each climatic variable while including phylogeny as a covariate in partial RDA models. Considering possible correlations among climatic variables, we conducted additional permutation tests for the marginal effects of climatic variables in a model including all climatic variables by setting the parameter by =’margin’ in anova.cca. This method tested the significance of each climatic variable while controlling for the effects of the others.

Traits correlated with climate variation were identified using a partial RDA, treating all 25 traits as response variables and the six climatic variables as predictors. Since only the first RDA-constrained axis was significant for each of the climate variables, we searched for candidate traits associated with environmental adaptation along that axis, selecting those with loadings exceeding ±2 standard deviations (SDs) from the mean loading. Trait-climatic variable associations were summarized by extracting, for each trait, the climatic variable with the largest absolute RDA loading. To test trait-climatic variable correlations, pairwise correlations between candidate traits and climatic variables were computed using Pearson’s correlation coefficients. Statistical significance was evaluated using a two-sided correlation test, and resulting *p*-values were adjusted for multiple testing using the false discovery rate (FDR) approach (*p*-values < 0.01).

## Results

Principal component analysis (PCA) of trait covariation in the 45 annual and perennial *Hordeum* accessions revealed that annual accessions occupied a larger portion of the multivariate trait space than perennial accessions (Fig. 2a, b). Grain size and weight, stem diameter, dry shoot biomass, and leaf size contributed most to PC1, which explained 38.5% of the overall trait variation and separated *H. vulgare* (wild and cultivated barley) from all other annual and perennial *H.* accessions (Figs. 2a,b, S1). All *H. vulgare* genotypes showed significantly greater values on PC1 than the other annual and perennial *Hordeum* accessions (Fig. 2c), indicating faster vegetative growth and larger organ sizes for leaves, stems, and grains. The remaining annual *Hordeum* accessions did not separate from perennial accessions in mean performance along PC1 (Fig. 2c).

**Fig. 2.**
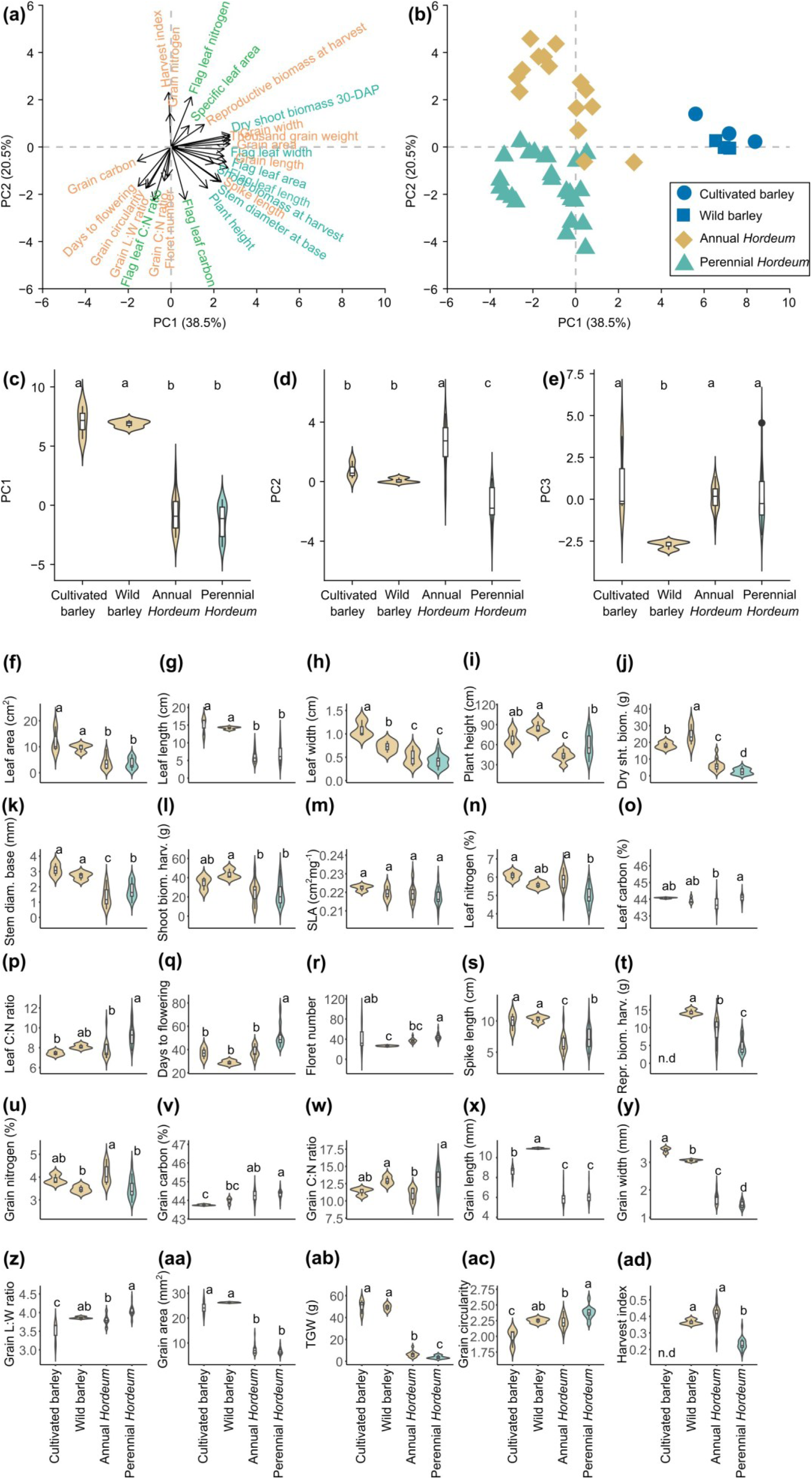
Principal component analysis (PCA) of 45 *Hordeum* accessions showing multivariate trait variation structured along the first two principal components. **(a)** PCA for 25 trait variables. The length of each black arrow from the origin indicates the strength with which that trait is represented in a given region of the principal component space. Traits are categorized into vegetative (dark green), leaf economic spectrum (light green), and reproductive (orange) traits. **(b)** Individual accession values are mapped to the same principal component space, with color and shape separating *H. vulgare* (blue, cultivated, n = 3; wild, n = 3), annual (brown, n = 15), and perennial accessions (teal, n = 24). **(c-e)** Violin plots show the distribution of each PC axis, with an embedded boxplot comparing median differences in principal components (PC1-PC3) between life history sub-groups. (**f-ad)** Violin plots show the distribution of each trait, with overlaid boxplots comparing the median values of annual (brown, cultivated, n = 3, wild, n =3, annual *Hordeum*, n = 15) and perennial (teal, n = 24) *Hordeum* accessions. Letters show the results of a post hoc test (Least Significant Difference, LSD) based on median pairwise comparisons (*p* < 0.05) between life history sub-categories. Different letters indicate significant differences between the sub-categories. Flag leaf C:N, Flag leaf carbon-to-nitrogen ratio; Grain L:W, Grain length-to-width ratio; Grain C:N, Grain carbon-to-nitrogen ratio. SLA, Specific leaf area; Shoot biom. harv., Shoot biomass at harvest; Dry sht. biom., Dry shoot biomass 30-days after planting (30-DAP); Repr. biom. harv., Reproductive biomass at harvest; Stem diam. base, Stem diameter at base; TGW, Thousand-grain weight; n.d, missing data.

PC2 (20.5% of variance) separated annual from perennial *Hordeum* accessions. Annual accessions had higher grain and leaf nitrogen concentrations, higher harvest index (Figs. 2a,b,n,u,ab, S1), whereas perennials exhibited higher grain and leaf carbon concentrations, C: N ratios, later flowering time, increased grain length-to-width ratio, and grain circularity (Figs. 2a,b,d,p,q,v,w,x,ac,z S1). Variation along PC3 (9.7% of variance) was primarily controlled by floret number and grain length, and distinguished wild from cultivated barley (Fig. 2e, S1). Cultivated barley showed higher values for floret number, primarily driven by the six-rowed barley cultivar Morex, whereas increased grain length was associated with wild barley (Figs. 2r,x, S1). In the phylogeny-corrected PCA, trait-accession covariance was largely comparable along PC1 and PC2 compared to the PCA uncorrected for phylogeny (Fig. S2a-d). However, PC3 primarily separated accessions according to the grain C:N ratio, which was strongly differentiated between annual and perennial *Hordeum* species (Fig. S2f).

We then tested whether trait covariation and thus trait syndromes consistently differ between annual and perennial *Hordeum* accessions by calculating pairwise Pearson’s correlation coefficients within the annual and perennial *Hordeum* subgroups separately and across all accessions (Figs. 3, S3). The annual accessions showed strong positive correlations of the vegetative traits, leaf size, plant height, and biomass with reproductive traits, i.e., grain size and weight. In the perennial accessions, these vegetative traits were not correlated with grain size or weight, but were correlated with spike length and floret number (Fig. 3).

**Fig. 3.**
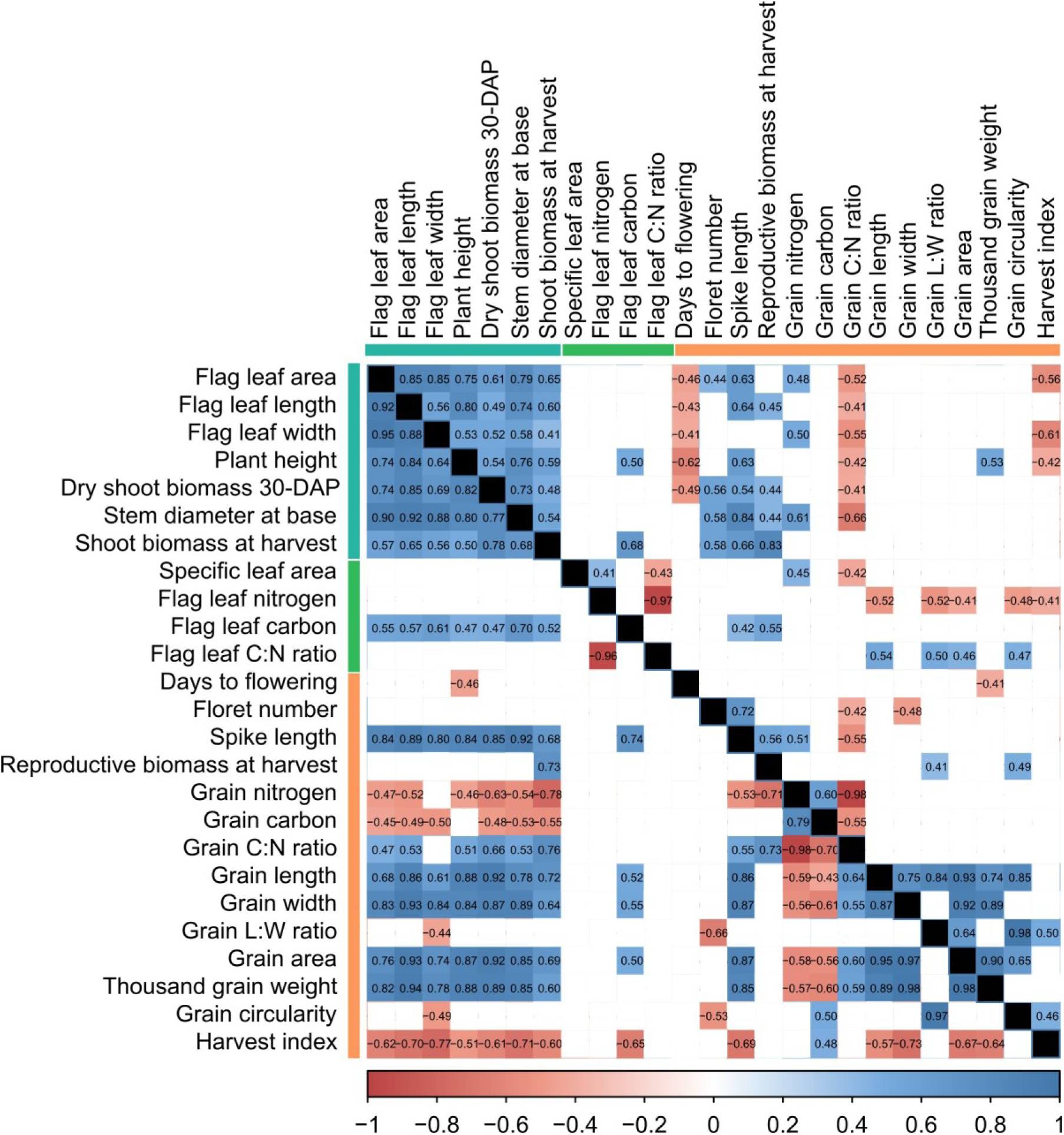
Pearson correlation matrix for trait variation within all annual accessions (lower diagonal, n = 21), and perennial accessions (upper diagonal, n = 24). Blue and red squares denote significant positive and negative correlations, respectively (*p* < 0.05, Bonferroni-adjusted). Traits are categorized into vegetative (green), leaf economic spectrum (LES, light green), and reproductive (orange) traits using colored bars along the top and left sides of the figure. Flag leaf C: N, Flag leaf carbon-to-nitrogen ratio; Grain C: N, Grain carbon-to-nitrogen ratio; Grain L: W, Grain length-to-width ratio; Dry shoot biomass 30-DAP, Dry shoot biomass 30-days after planting.

Furthermore, the vegetative traits, flag leaf length, plant height, shoot biomass at 30-days after planting (DAP), and at harvest, were negatively correlated with grain C and N contents in the annual subgroup, reflecting a trade-off between investing in vegetative versus reproductive growth. In perennials, the vegetative traits, flag leaf area, SLA, and stem diameter were positively correlated with grain nitrogen, indicating coordinated variation between vegetative growth and reproductive nutrient allocation. N and C contents in leaf and grain were not correlated in either the annual or the perennial subgroups, indicating that nutrient concentrations in vegetative tissues do not correspond to nutrient allocation to reproductive tissues across life histories.

The leaf economic traits, SLA, leaf N, and C contents, were only correlated in the perennial subgroup. SLA was positively correlated with leaf and grain N and negatively correlated with leaf and grain C: N ratios in the perennial but not the annual subgroups. SLA was not correlated with dry shoot biomass at 30-DAP, a proxy for vegetative growth rate, in either the annual or the perennial subgroups (Fig. 3).

Because overall trait variation was influenced by both life history and phylogeny, we next quantified the effects of life history on individual traits after accounting for phylogeny. Life history significantly contributed to variation in 14 of the 25 measured traits (Fig. 4). Six traits, harvest index, flag leaf and grain nitrogen, grain width, TGW, and reproductive biomass at harvest, were significantly higher in annual compared to perennial *Hordeum* accessions (Fig. 4). By contrast, values for grain C:N ratio, grain length:width ratio, and circularity, days to flowering, plant height, stem diameter, flag leaf C content, and C:N ratio, were higher in the perennial *Hordeum* species. Life history had the strongest effects on harvest index (51%), grain N content (26%), grain C:N ratio (25%), flag leaf C (24%), and days to flowering (22%; Table S6). Although annual and perennial accessions differed significantly on average for these traits, their trait distributions overlapped, indicating substantial within-group variation (Fig. S4a-y).

**Fig. 4.**
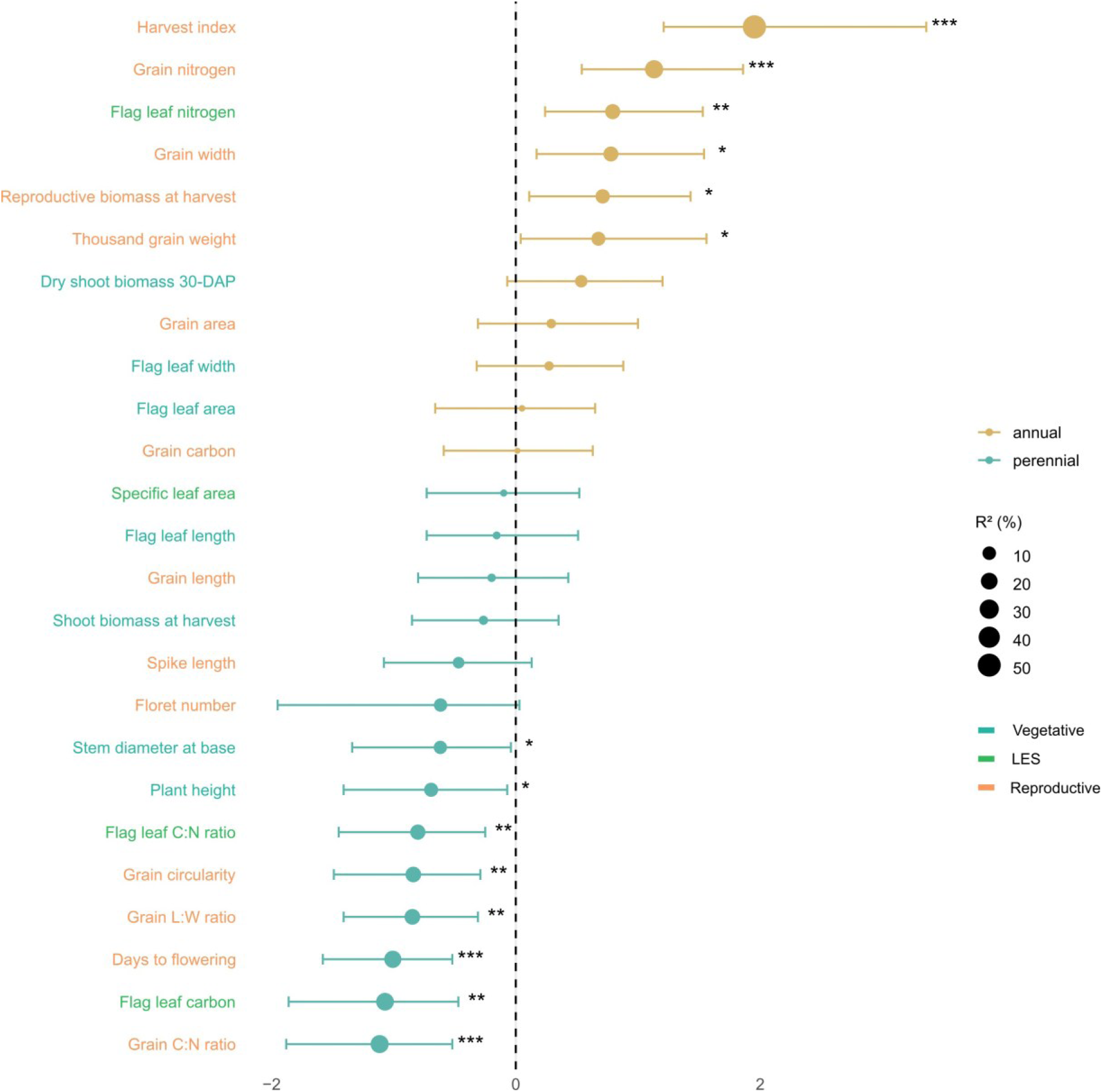
Forest plot showing differences in phylogenetically corrected trait values between annual and perennial species. Traits and their respective effect sizes between the life history groups are categorized into vegetative (green), leaf economic spectrum (LES, light green), and reproductive (orange) traits. Points represent effect sizes (annual vs. perennial), and traits are ordered by the magnitude of effect size relative to life history strategies and are expressed as percentages derived from *R*^2^. Effect sizes are reported as Cohen’s *d* with 95% confidence intervals. Significance levels: *, *p* < 0.05; **, *p* < 0.01; ***, *p* < 0.001. Four accessions, *H. euclaston* (NGB90342), *H. bogdanii* (NGB90046), and *H. chilense* (GRA972, NGB90118) were removed from this analysis because they were not included in the phylogenetic tree. Flag leaf C: N, Flag leaf carbon-to-nitrogen ratio; Grain C: N, Grain carbon-to-nitrogen ratio; Grain L: W, Grain length-to-width ratio; Dry shoot biomass 30-DAP, Dry shoot biomass 30-days after planting.

Taken together, annual *Hordeum* accessions occupied a larger trait space, largely driven by the distinctive performance of annual wild and cultivated barley. Trait covariation differed between life histories: in annuals, vegetative growth was positively associated with grain size and weight but negatively with grain nutrient content, whereas in perennials, vegetative traits were positively linked to floret number and grain N.

Annual and perennial *Hordeum* accessions differed in traits associated with allocation to reproductive output (harvest index), grain nutrient composition (N content, C:N ratio), leaf C investment, and flowering time. Despite significant average trait differences, overlapping trait distributions indicate phenotypic continuity rather than discrete separation between the two life-history groups.

### Association between traits and bioclimatic variables

Life history is often considered to result from adaptations to different environments (Stearns, 1992). To investigate the environmental correlates of trait variation within and between life-history groups, we tested associations between trait variation and bioclimatic variables of the accessions’ native habitats. First, we selected six uncorrelated bioclimatic variables that together represented the variation across all 19 original variables (Fig. S5b,c). We then used redundancy analyses (RDA) to relate trait variation to climatic conditions at the accessions’ origins, using the six selected climate variables. We found that a large part (33%) of the overall trait variation was explained by the combination of the six bioclimatic variables (precipitation levels and temperature) in the simple RDA model, which was not corrected for phylogeny (Table S7, Fig. 5d). After correcting for phylogeny in a partial RDA, the bioclimatic variables still explained a significant proportion of trait variation (15%, Table S7). Annuals and perennials overlapped in trait–climate space, indicating broadly conserved relationships between traits and climatic variables across life histories. Perennials exhibited greater dispersion along both axes, suggesting a wider ecological breadth or increased variability in their trait–environment associations. However, annuals and perennials were partially separated along the primary climatic gradient (RDA1), with perennials associated with conditions represented by positive axis values and annuals clustering toward negative to intermediate values, indicating divergent trait–environment syndromes between life histories. Since the combination of six selected variables explained a significant proportion of the trait variation in the RDA models, we further investigated the effects of the individual climatic variables on overall trait variation. By sequentially fitting one variable at a time in a simple RDA, we found that four climatic variables, isothermality (Bio3_T), temperature seasonality (Bio4_T), mean temperature of the driest quarter (Bio9_T), and precipitation of the coldest quarter (Bio19_P), were significantly associated with trait variation (*p* < 0.05, Table 1a, Fig. 5a,b). Using partial RDA models conditioned on phylogeny, we confirmed that Bio9_T (9%, *p* = 0.004) and the mean temperature of the wettest quarter, Bio8_T (7%, *p* = 0.014, Fig. 5a,c, Table 1b), explained significant proportions of trait variation. Both Bio9_T and Bio8_T remained significantly associated with overall trait variation even after accounting for the marginal effects of other climatic variables, explaining 6% and 5% of trait variation, respectively (Table S8). The RDA analysis also demonstrated that the mean temperature of the driest quarter (Bio9_T) and the mean temperature of the wettest quarter (Bio8_T) were associated with the direction along which annuals and perennials differed in trait expression (Fig. 5b). Such contrasting covariation suggested that temperature seasonality differentially structured trait variation across annual and perennial *Hordeum* species.

**Fig. 5.**
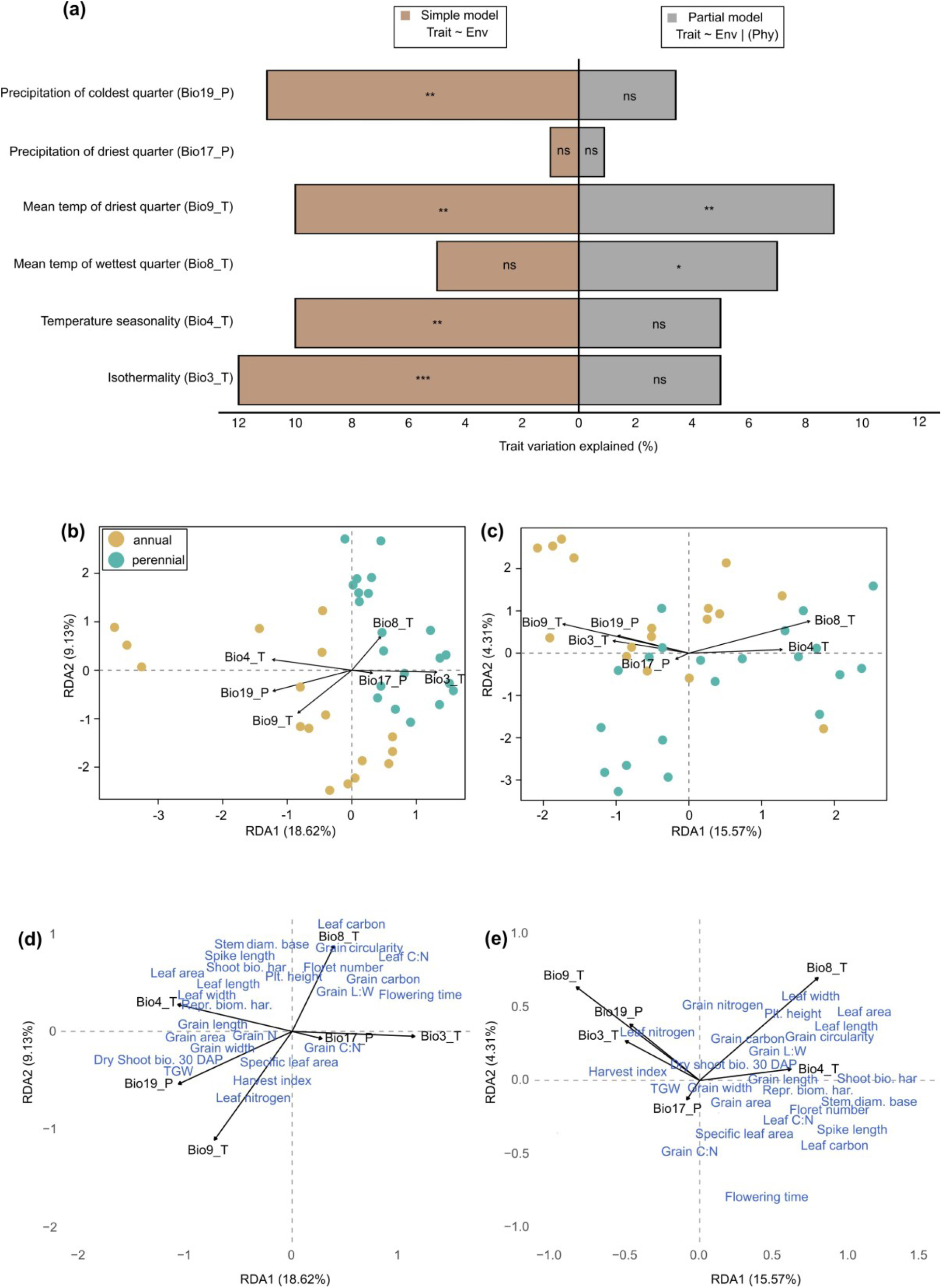
Partitioning of trait variance explained by climatic variables using redundancy analysis (RDA) **(a)** Beige bars represent the percentage of trait variation explained by individual climatic variables without phylogenetic correction (simple model), while grey bars show values after accounting for phylogenetic relatedness (partial model). Significance levels: ns, *p* > 0.1; *, *p* < 0.05; **, *p* < 0.01; ***, *p* < 0.001. **(b-e)** Biplot of trait-climatic variable associations for simple (b,d) and partial RDA (c,e). Solid black arrows (black labels) represent climatic variables, and blue text represents traits in **d-e.** For visualization, accession-climatic **(b,c)** and trait-climatic variable associations **(d,e)** are shown separately. However, in b and c, individual accession trait values are mapped to the same principal component space, with color separating annual accessions (brown, n = 17) and perennial accessions (teal, n = 21). Abbreviations: Bio3_T, Isothermality; Bio4_T, Temperature seasonality; Bio8_T, Mean temperature of wettest quarter; Bio9_T, Mean temperature of driest quarter; Bio17_P, Precipitation of driest quarter; Bio19_P, Precipitation of coldest quarter.

**Table 1:** Effects of individual climatic variables on trait variation as estimated by redundancy analysis (RDA)

| <sup>a</sup>Model type | Climatic variable | *F*-value | Var | Var (%) | *p*-value |
| --- | --- | --- | --- | --- | --- |
| Individual effect of climatic variables | Bio3_T | 5.20 | 2.94 | 11.77 | 0.0004*** |
|  | Bio19_P | 4.86 | 2.77 | 11.07 | 0.0024** |
|  | Bio4_T | 4.56 | 2.62 | 10.47 | 0.0012** |
|  | Bio9_T | 4.20 | 2.43 | 9.71 | 0.0014** |
|  | Bio8_T | 2.13 | 1.29 | 5.18 | 0.0636 |
|  | Bio17_P | 0.40 | 0.25 | 1.01 | 0.9114 |
| <sup>b</sup>Individual effect of climatic variables conditioned on phylogeny | Bio9_T | 3.46 | 1.33 | 8.76 | 0.0044** |
|  | Bio8_T | 2.80 | 1.10 | 7.22 | 0.0144* |
|  | Bio4_T | 2.01 | 0.80 | 5.28 | 0.0668 |
|  | Bio3_T | 1.83 | 0.73 | 4.84 | 0.0896 |
|  | Bio19_P | 1.27 | 0.52 | 3.42 | 0.2515 |
|  | Bio17_P | 0.35 | 0.15 | 0.96 | 0.9540 |

Individual climatic variables are arranged according to explained trait variation (highest to lowest). Permutation tests computed *p*-values based on 5,000 permutations. Significance levels: $p > 0.1$ ; \*, $p < 0.05$ ; \*\*, $p < 0.01$ ; \*\*\*, $p < 0.001$ . Abbreviations: Bio3\_T, Isothermality; Bio4\_T, Temperature seasonality; Bio8\_T, Mean temperature of wettest quarter; Bio9\_T, Mean temperature of driest quarter; Bio17\_P, Precipitation of driest quarter; Bio19\_P, Precipitation of coldest quarter. Var: Variance

We next examined which specific traits were associated with these climatic drivers. To this end, we fitted a partial redundancy analysis (RDA) model that accounted for phylogenetic relatedness, thereby identifying trait–environment associations that were consistent across the phylogeny. This analysis yielded a single significant constrained axis (Table S9), along which one candidate trait, flag leaf N concentration (%), was significantly correlated with the mean temperature of the driest quarter (Bio9_T, 0.51, *p* = 0.0038, Table S10, Figs. 5e, S6). This suggested that the elevated flag leaf N concentration characteristic of annual species is associated with warmer dry-season conditions.

In summary, the observed patterns suggested that while annual and perennial species share a conserved trait–climate relationships, life history drives differentiation in how these traits are expressed along climatic gradients. Mean temperatures of the driest (Bio9_T) and wettest (Bio8_T) quarters were aligned with the axis, differentiating annual and perennial *Hordeum* species in trait composition, indicating that temperature seasonality structures trait variation, with annual traits associated with warmer dry seasons and perennial traits associated with warmer wet seasons. Specifically, high flag leaf N was positively associated with Bio9_T, suggesting that environments experiencing strong terminal stress may have selected for elevated flag leaf N, a characteristic of fast-cycling annual *Hordeum* species. The broader dispersion of perennials in the trait-climate relationship indicated greater ecological flexibility, consistent with adaptation to a wider range of environmental conditions.

## Discussion

### Wild and cultivated barley show strong trait divergence from all other *Hordeum* species

To identify genetic variation in traits and trait combinations that distinguish annual and perennial *Hordeum* accessions, we applied a phylogenetic framework to 25 functional traits measured in a common-garden experiment at a single location over three years. Annual accessions occupied a larger trait space, driven by the distinct trait profile of the annual *H. vulgare* (barley), which exhibited higher growth rates and larger organs (leaves, stems, and grains) than other *Hordeum* accessions. Growth and organ size variation within the *Hordeum* clade were thus primarily controlled by phylogeny rather than life history, with barley displaying fast growth rates, in contrast to the conservative growth of other wild annual and perennial *Hordeum* accessions. Interestingly, this separation was independent of domestication, as wild barley clustered with domesticated barley cultivars, supporting the idea that high growth rates and relatively large grains may have predisposed it for domestication (Weide et al., 2018; Romero et al., 2025). Similarly, a comprehensive study of the evolution of growth rates in wild, landrace, and improved accessions of 19 herbaceous crops demonstrated that domestication and improvement breeding had no effect on growth rates in barley and other cereal crops (Gómez-Fernández et al., 2022). Compared to variation between wild *Hordeum* species, the trait differences introduced by domestication and breeding were thus relatively minor. Aside from grain shape, key domestication traits, such as leaf, stem, and spike sizes, and grain weight, showed little divergence between wild and cultivated barley. Indeed, unlike some other crops, the visible phenotype of domesticated barley remains largely similar to its wild form, with spike rachis brittleness being the only well-characterized trait showing clear dimorphism between wild and domesticated subgroups (Abbo et al., 2014; Pourkheirandish et al., 2015). This limited phenotypic divergence is consistent with genomic evidence indicating a protracted domestication process and ongoing gene flow between wild and cultivated barley (Pankin et al., 2018; Guo et al., 2025).

### Annuals and perennials differ in resource allocation but show extensive trait overlap

We identified significant trait differences between annual and perennial *Hordeum* accessions along PC2 (Fig. 2a, b), primarily driven by variation in flowering time, dry shoot biomass, floret number, grain circularity, harvest index, leaf N and C, and grain N. However, a strong overlap in trait values among annual and perennial *Hordeum* accessions indicated that these did not consistently separate by life-history strategy. These patterns align with previous findings that phylogenetic classification often explains a larger proportion of trait variation than life-history type (Mazer, 1989; Vico et al., 2016; Herron et al., 2021), suggesting that historical and lineage-specific constraints shape functional trait expression within closely related annual and perennial species (Silvertown & Dodd, 1996).

Vegetative and reproductive biomass were positively correlated in both life history groups, but strategies differed: in annual *Hordeum* accessions, vegetative biomass was correlated with grain size traits, whereas in perennials, it was correlated with floret numbers. This suggested that the annuals favored investment in grain size, producing wider, heavier grains to increase the likelihood of seedling establishment (Leger et al., 2019; Quigley et al., 2023). By contrast, perennials produced more, but smaller, thinner, and lighter grains than annual accessions, thereby likely facilitating dispersal (Vico et al., 2016; Herron et al., 2021; Gao et al., 2024). This contrasts with findings in the legume families Lathyrus, Phaseolus, and Vicia, where seed size was not consistently different between annuals and perennials, but was clade-specific (Herron et al., 2021). On the other hand, a study in the genus Astragalus demonstrated that perennials tended to have more flowers and ovules than annuals, consistent with the higher floret numbers observed in perennial *Hordeum* accessions (Sinjushin et al., 2024).

The leaf economics spectrum describes a major axis of global plant functional trait variation that differentiates annual resource-acquisitive from perennial resource-conservative strategies (Garnier et al., 1997; Segrestin et al., 2020; Ning et al., 2022). Perennial *Hordeum* accessions exhibited slower growth, delayed flowering, and carbon-rich leaves, reflecting a resource-conservative strategy consistent with the leaf economics spectrum and adaptation to stress-prone, resource-poor environments (Grime, 1977; Coley et al., 1985; Wilson & Thompson, 1989; Wright et al., 2004). By contrast, annual *Hordeum* accessions invested in more leaf N than perennial accessions, which has been linked to higher photosynthetic rates (Field & Mooney, 1986; Reich et al., 1997; Niinemets, 1999; Wright et al., 2001).

SLA is a key leaf economics trait that correlates with whole-plant growth and has been reported to differ between fast-growing annual and resource-conservative perennial species (Liu et al., 2017; Ning et al., 2022). However, SLA was not significantly different between annual and perennial *Hordeum* accessions (Fig. 3) and was not correlated with biomass growth or plant longevity (Fig. S3). This result contrasts with previous studies conducted at broader taxonomic scales (Garnier, 1992; Garnier et al., 1997), suggesting that leaf-level trade-offs may not hold within closely related species where shared evolutionary history constrains trait differentiation. A study across 28 species of the diverse herbaceous genus *Helianthus* (sunflowers) also demonstrated that leaf-level trait variation was not related to life span, suggesting that leaf economics patterns do not reflect universal physiological trade-offs at small evolutionary scales (Mason & Donovan, 2015).

Faster growth and higher harvest index in annual versus perennial *Hordeum* accessions indicate a strategy that prioritizes reproduction over extended vegetative growth. This aligns with a recent study in the grass subfamily Pooideae, which suggests that high relative growth rates and preferential allocation to above-ground structures were key evolutionary precursors to the transition from perennial to annual life histories (Lindberg et al., 2020).

Lower growth rates in perennial accessions were compensated by a longer growth period during the season, resulting in comparable shoot biomass at harvest between annuals and perennials (Fig. 4). Successive waves of tiller production persisting until the termination of the experiments supported the notion that slow-growing species can accumulate greater total biomass over time (Midgley & Bond, 1991; Westoby et al., 2002). We speculate that perennial *Hordeum* accessions also allocate more resources to below-ground root biomass over time to support future above-ground growth, a hypothesis warranting detailed investigation of root traits.

### Temperature variation drives trait variation between annual and perennial *Hordeum* accessions

Life cycle variation has been interpreted as an adaptation to different climatic conditions, in which annuals and perennials manage trade-offs between current survival and future reproduction differently (Poppenwimer et al., 2023). While annuals focus on rapid reproduction within one season, perennials focus on long-term survival across multiple seasons (Volis et al., 2002; Sherrard & Maherali, 2006; Jaikumar et al., 2012; Segrestin et al., 2018; Gao et al., 2024).

We demonstrated that annual and perennial *Hordeum* species were separated in RDA space defined by bioclimatic variables, suggesting that the two life histories’ distinct phenotypic trait syndromes are associated with climatic gradients. In our results, trait variation separating annual and perennial accessions covaried with temperature, with annual traits positively linked to temperature in the driest quarter and perennial traits positively linked to temperature in the wettest quarter. This demonstrated that specific seasonal climate windows, rather than annual averages, determine trait variation in *Hordeum*. This finding aligns with previous studies showing that the prevalence of annual species in different taxonomic clades is driven by temperature and precipitation in the driest quarter rather than by annual means (Fiz et al., 2002; Evans et al., 2005; Monroe et al., 2019; Poppenwimer et al., 2023). Unlike many studies emphasizing precipitation as the primary constraint, annual traits in *Hordeum* were associated with temperature, suggesting that thermal stress during the dry season represents a stronger ecological filter than water availability. This is consistent with the distribution of annual and perennial *Hordeum* species, as annuals occur in warmer regions, particularly during summer, than perennials, whereas precipitation does not differ markedly between their habitats (Hellwig et al., 2025). Leaf N emerged as a key trait distinguishing annual and perennial *Hordeum* species, with annuals consistently exhibiting higher leaf N concentrations than perennials. High leaf N has been linked to higher photosynthetic capacity, faster growth, and accelerated development, especially in fast-growing annual species (Mooney, 1986; Field & Evans, 1989). Leaf N variation was positively correlated with temperature in the driest quarter, suggesting that higher photosynthetic capacity may have been selected in annual *Hordeum* to drive growth and development under conditions of terminal stress (Griffin-Nolan et al., 2026). Perennial traits showed a significant correlation with temperature in the wettest quarter, suggesting that trait variation in perennial *Hordeum* species is linked to climatic conditions during the main growth period, when water availability is high, and temperature influences growth, survival, and resource allocation in long-lived plants (Münzbergová et al., 2024; Journé et al., 2025). Because perennials invest in long-lived structures, trait variation may be particularly sensitive to climatic conditions during the main growth season, when water availability is not limiting. Similar interactions between temperature and moisture have been shown to influence functional trait variation and genetic adaptation in perennial plants in which growth conditions during the favourable season strongly determine long-term performance (Journé et al., 2025).

## Conclusion

In conclusion, annual *H. vulgare* stands out within the *Hordeum* clade for its high growth rates and large organs, a phylogenetically driven, acquisitive strategy that predates domestication, while other annual and perennial species show rather conservative growth traits, highlighting that domestication and breeding introduced relatively minor changes to growth and organ size traits compared to trait variation between *Hordeum* species. Consequently, growth and overall plant productivity are largely determined by phylogeny rather than life history within the *Hordeum* clade.

Annual and perennial *Hordeum* accessions primarily differ in resource allocation: annuals invest in higher leaf N and larger grains, while perennials produce carbon-rich leaves and smaller, more numerous grains. Yet trait variation strongly overlaps between annual and perennial *Hordeum* accessions, indicating that, in addition to life-history differences, both phylogenetic history and climatic conditions at the accession’s origin contribute to shaping functional variation within and between annual and perennial species. Trait divergence between annual and perennial *Hordeum* reflects adaptation to different seasonal thermal environments, with annual traits linked to temperature in the driest quarter and perennial traits linked to temperature in the wettest quarter. This suggests that annual trait variation is primarily shaped by unfavorable conditions at the end of the growing season that prevent survival and favor persistence through seeds. In contrast, perennial trait variation is influenced by climatic conditions during the main growth phase, which determine long-term growth performance and survival. The overlap in trait–climate space suggests that annuals and perennials can occur under similar climatic regimes, but trait variation is shaped by end-of-season stress in annuals and by conditions during peak growth in perennials.

## Acknowledgements

The authors thank Maria Graf (CMML, Heinrich Heine University Düsseldorf) for technical support in analyzing leaf nitrogen and carbon. We thank Andreas Fischbach and the gardeners at the HHU Botanical Garden in Düsseldorf for their support of the common garden experiments. This work was funded by the European Research Council (ERC) under the European Union’s Horizon Europe research and innovation program (PERLIFE, No. 101002085) and the Deutsche Forschungsgemeinschaft (DFG) under Germany’s Excellence Strategy (EXC-2048 – Project ID: 390686111 and the Collaborative Research Centre/Transregio (TRR 341, Project ID: 456082119)

## Author contributions

MA: Conceptualization, Investigation, Methodology, Software, Formal analysis, Visualization, Writing – original draft, Data curation, Project administration. TH: assisted in Formal analysis, Methodology, Writing – review & editing. EBH: Methodology and data management. RS: Investigation. ND: Investigation. PW: Carbon and nitrogen measurement. AB: Methodology, review & editing. MvK: Conceptualization, Funding acquisition, Resources, Writing – review & editing, Project administration, Supervision.

## Data availability

The data that support the findings of this study are openly available at DataPLANT (Weil et al., 2023), our FAIR data publication can be found under doi: https://doi.org/10.60534/cvcp1-dfd77.

## Supporting Information – Supplemental Tables

**Table S1:** List of *Hordeum* accessions used in the study.

**Table S2:** Air temperature (°C) with mean standard deviation (±SD) for day, night, and overall, by month and year.

**Table S3:** Description of phenotypic traits, functional importance, references, and the years in which traits were evaluated.

**Table S4:** Traits BLUP-adjusted means for all accessions used in the study.

**Table S5:** All environmental variables retrieved from the WorldClim2 database and used in spatial niche modelling to identify the variables that best describe the niches of the different *Hordeum* accessions.

**Table S6:** Comparison of traits between annual and perennial species after accounting for phylogeny.

**Table S7:** Trait-environment associations.

**Table S8:** Effects of individual climatic variables estimated by RDA models.

**Table S9:** Trait-environment associations for all RDA axes.

**Table S10:** Candidate trait-environment associations detected only on the first significant constrained RDA axis after accounting for phylogenetic relatedness.

## Supporting Information - Supplemental figures

**Fig. S1:** Heatmap of PCA for 45 *Hordeum* accessions, showing the trait space defined by PC1-PC3.

**Fig. S2:** Phylogeny corrected PCA of 41 *Hordeum* accessions, showing the trait space defined by principal components.

**Fig. S3:** Pearson correlation matrix of the full dataset, including annual (n = 21) and perennial (n = 24) *Hordeum* accessions.

**Fig. S4:** Boxplot showing mean trait values per accession, averaged across the 2021-2023 growing seasons (n = 8-12).

**Fig. S5:** Procedure for generating synthetic environmental variables based on the methods described by Chang et al. (2022).

**Fig. S6:** Associations between traits and climatic variables along the first significant constrained RDA axis after accounting for phylogenetic relatedness.

