## Supplementary Figures and Tables for "Phylogeny, thermal seasonality, and life history shape trait variation in annual and perennial barley relatives"

### Supplemental Figures

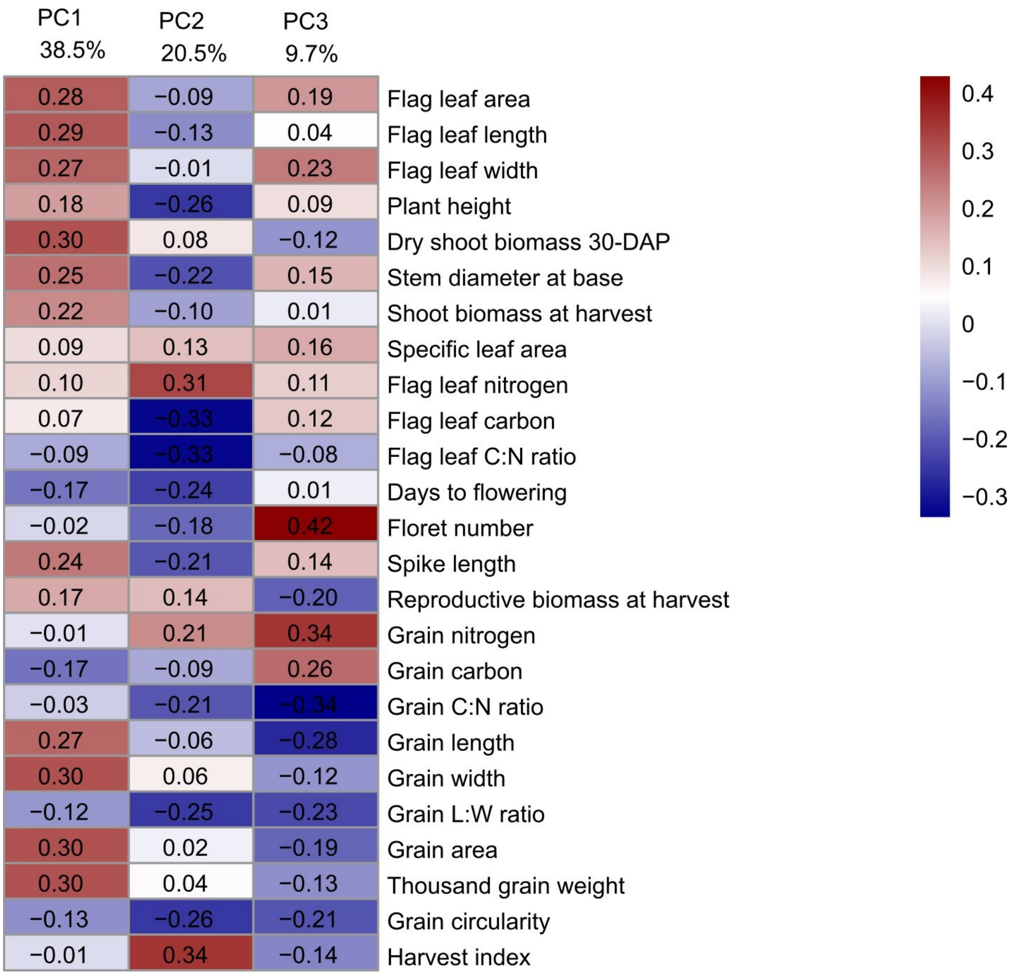


**Fig. S1: Heatmap of principal component analysis (PCA) for 45 Hordeum accessions, illustrating the distribution of trait variation across the multivariate trait space defined by the first three principal components (PC1–PC3).**

The heatmap represents the contribution of traits to principal components, summarized to indicate the strength and direction of the associations. Red and blue colors represent positive and negative loadings, respectively.

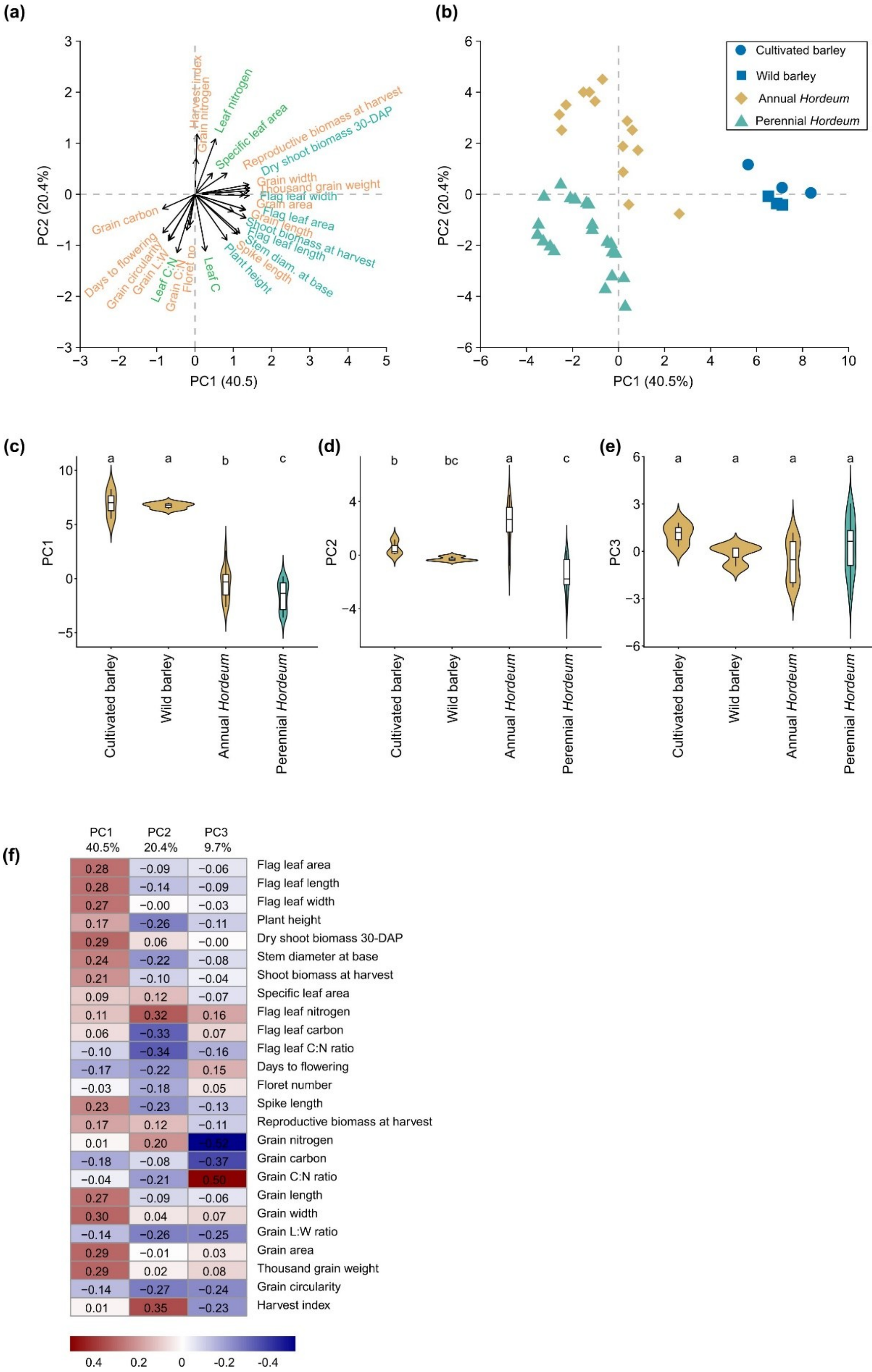


**Fig. S2: Phylogeny corrected PCA of 41 *Hordeum* accessions, showing the trait space defined by the three first principal components.**

**(a)** PCA showing 25 trait variables. The length of each black arrow from the origin indicates the strength with which that trait is represented in a given region of the principal component space Traits are categorized into vegetative (dark green), leaf economic spectrum, LES (light green), and reproductive (orange). **(b)** Individual accession values are mapped to the same principal component space, with color and shape separating annual accessions (brown, n = 20), perennial accessions (teal, n = 21). **(c-e)** Violin plots show the distribution of each PC axis, with an embedded boxplot comparing median differences in principal components (PC1-PC3) between life history sub-groups. Letters show the results of a post hoc test (Least Significant Difference, LSD) based on median pairwise comparisons ($p < 0.05$) between sub-groups. Different letters indicate significant differences between the groups. Leaf C:N, Flag leaf carbon-to-nitrogen ratio; Leaf C, Flag leaf carbon; Grain L:W, Grain length-to-width ratio; Grain C:N, Grain carbon-to-nitrogen ratio; Floret no., Floret number; Stem diam. at base, Stem diameter at base. **(f)** Heatmap representing the contribution of traits to principal components, summarized to indicate the strength and direction of the associations. Red and blue colors represent positive and negative loadings, respectively.

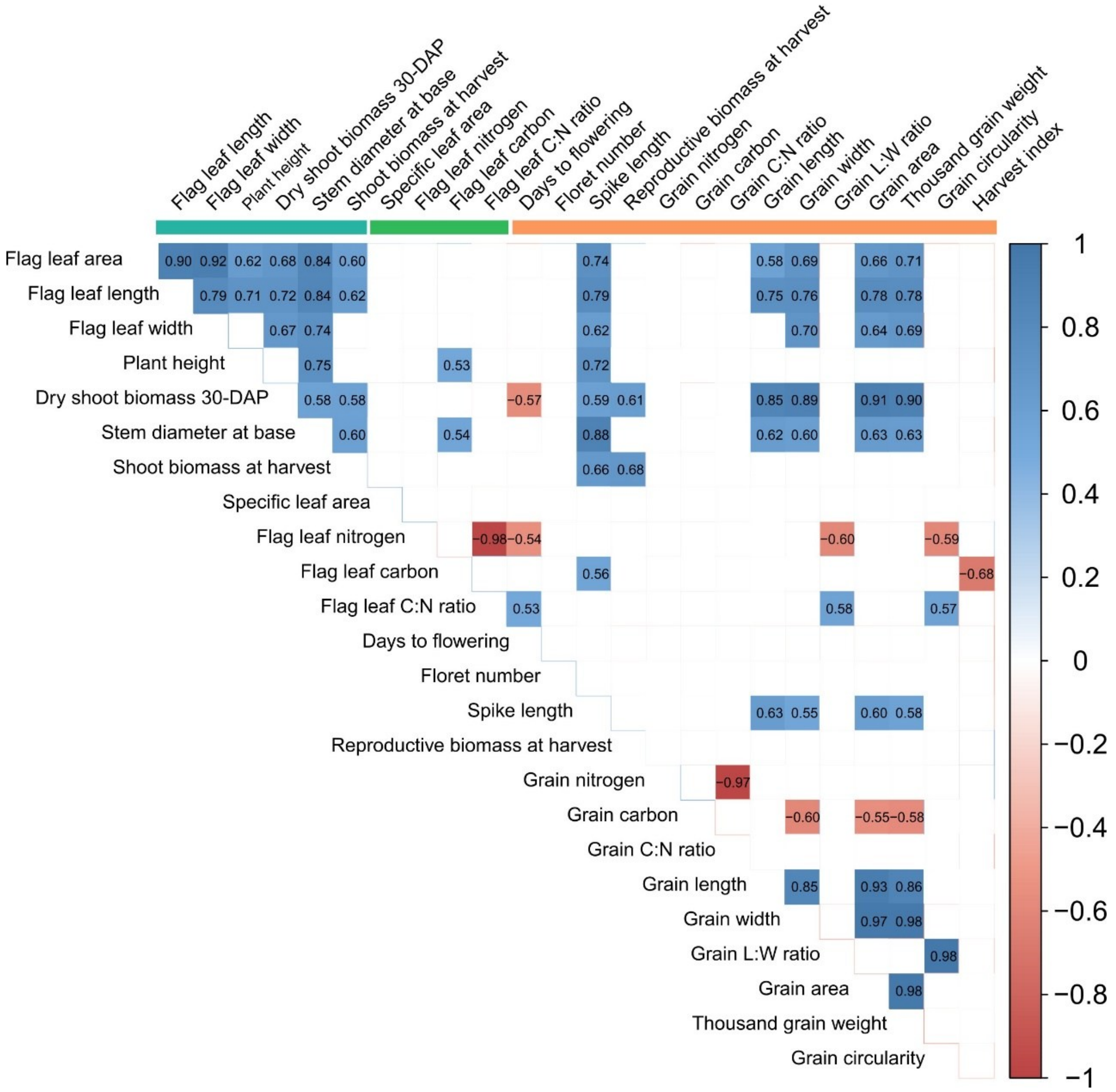


**Fig. S3: Pearson correlation matrix of the full dataset, including annual (n = 21) and perennial (n = 24) *Hordeum* accessions.**

Blue and red squares denote significant positive and negative correlations, respectively ($p < 0.05$, Bonferroni-adjusted). The traits are categorized into vegetative (green), LES (light green), and reproductive (orange) traits using colored bars along the top of the figure.

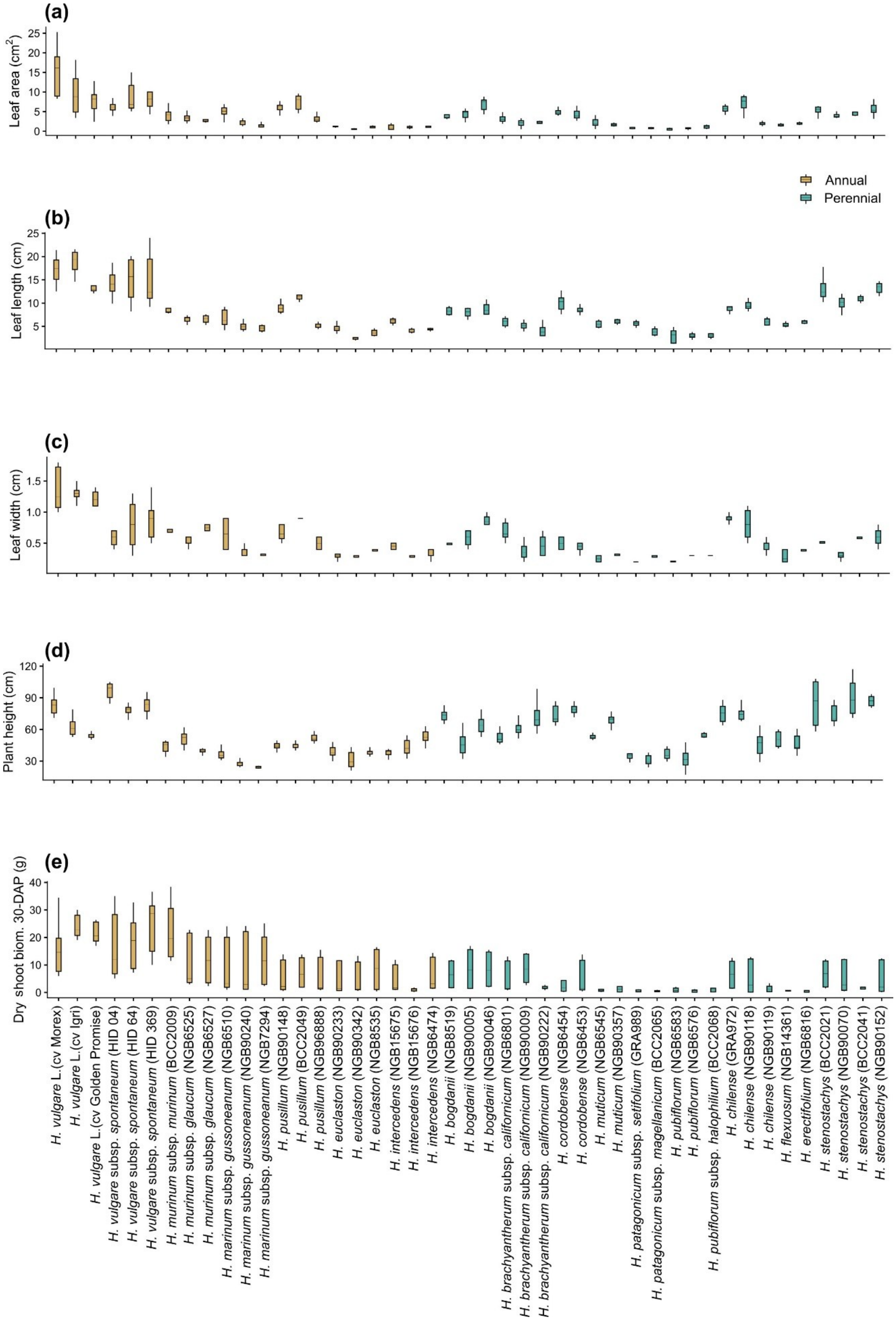

(a)
Leaf area (cm$^2$)
(b)
Leaf length (cm)
Annual
Perennial
(c)
Leaf width (cm)
(d)
Plant height (cm)
(e)
Dry shoot biom. 30-DAP (g)
H. vulgare L.(cv Morex)
H. vulgare L.(cv Igri)
H. vulgare L.(cv Golden Promise)
H. vulgare subsp. spontaneum (HID 04)
H. vulgare subsp. spontaneum (HID 64)
H. vulgare subsp. spontaneum (HID 369)
H. murinum subsp. murinum (BCC2009)
H. murinum subsp. glaucum (NGB6525)
H. murinum subsp. glaucum (NGB6527)
H. marinum subsp. gussoneanum (NGB6510)
H. marinum subsp. gussoneanum (NGB90240)
H. marinum subsp. gussoneanum (NGB7294)
H. pusillum (NGB90148)
H. pusillum (BCC2049)
H. pusillum (NGB96888)
H. euclaston (NGB90233)
H. euclaston (NGB90342)
H. euclaston (NGB8535)
H. intercedens (NGB15675)
H. intercedens (NGB15676)
H. intercedens (NGB6474)
H. bogdanii (NGB8519)
H. bogdanii (NGB90005)
H. bogdanii (NGB90046)
H. brachyantherum subsp. californicum (NGB6801)
H. brachyantherum subsp. californicum (NGB90009)
H. brachyantherum subsp. californicum (NGB90222)
H. cordobense (NGB6454)
H. cordobense (NGB6453)
H. muticum (NGB6545)
H. muticum (NGB90357)
H. patagonicum subsp. setifolium (GRA989)
H. patagonicum subsp. magellanicum (BCC2065)
H. pubiflorum (NGB6583)
H. pubiflorum (NGB6576)
H. pubiflorum subsp. halophilium (BCC2068)
H. chilense (GRA972)
H. chilense (NGB90118)
H. chilense (NGB90119)
H. flexuosum (NGB14361)
H. erectifolium (NGB6816)
H. stenostachys (BCC2021)
H. stenostachys (NGB90070)
H. stenostachys (BCC2041)
H. stenostachys (NGB90152)

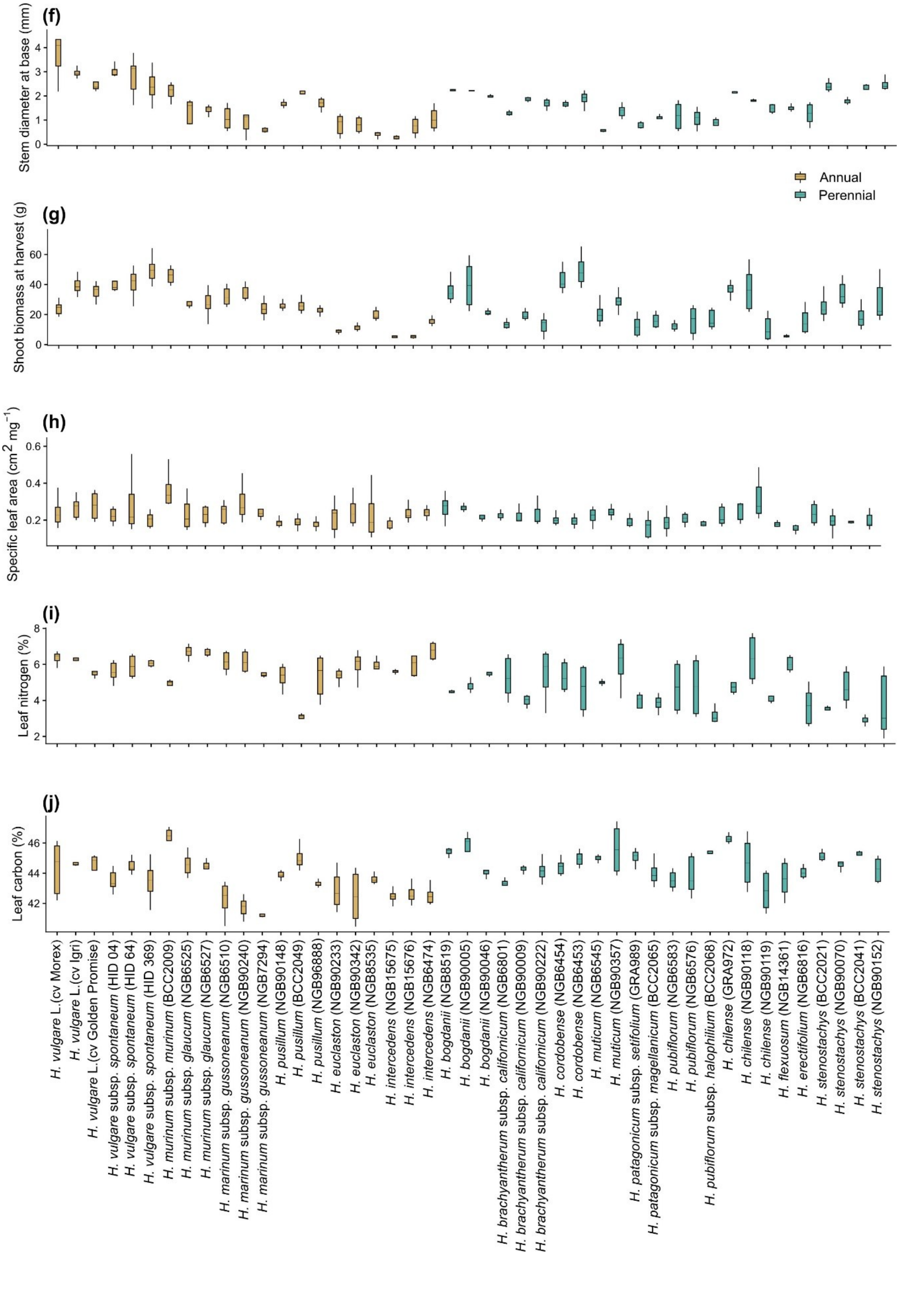

(f)
Stem diameter at base (mm)
Annual
Perennial
(g)
Shoot biomass at harvest (g)
(h)
Specific leaf area (cm2 mg−1)
(i)
Leaf nitrogen (%)
(j)
Leaf carbon (%)
H. vulgare L.(cv Morex)
H. vulgare L.(cv Igri)
H. vulgare L.(cv Golden Promise)
H. vulgare subsp. spontaneum (HID 04)
H. vulgare subsp. spontaneum (HID 64)
H. vulgare subsp. spontaneum (HID 369)
H. murinum subsp. murinum (BCC2009)
H. murinum subsp. glaucum (NGB6525)
H. murinum subsp. glaucum (NGB6527)
H. marinum subsp. gussoneanum (NGB6510)
H. marinum subsp. gussoneanum (NGB90240)
H. marinum subsp. gussoneanum (NGB7294)
H. pusillum (NGB90148)
H. pusillum (BCC2049)
H. pusillum (NGB96888)
H. euclaston (NGB90233)
H. euclaston (NGB90342)
H. euclaston (NGB8535)
H. intercedens (NGB15675)
H. intercedens (NGB15676)
H. intercedens (NGB6474)
H. bogdanii (NGB8519)
H. bogdanii (NGB90005)
H. bogdanii (NGB90046)
H. brachyantherum subsp. californicum (NGB6801)
H. brachyantherum subsp. californicum (NGB90009)
H. brachyantherum subsp. californicum (NGB90222)
H. cordobense (NGB6454)
H. cordobense (NGB6453)
H. muticum (NGB6545)
H. muticum (NGB90357)
H. patagonicum subsp. setifolium (GRA989)
H. patagonicum subsp. magellanicum (BCC2065)
H. pubiflorum (NGB6583)
H. pubiflorum (NGB6576)
H. pubiflorum subsp. halophilium (BCC2068)
H. chilense (GRA972)
H. chilense (NGB90118)
H. chilense (NGB90119)
H. flexuosum (NGB14361)
H. erectifolium (NGB6816)
H. stenostachys (BCC2021)
H. stenostachys (NGB90070)
H. stenostachys (BCC2041)
H. stenostachys (NGB90152)

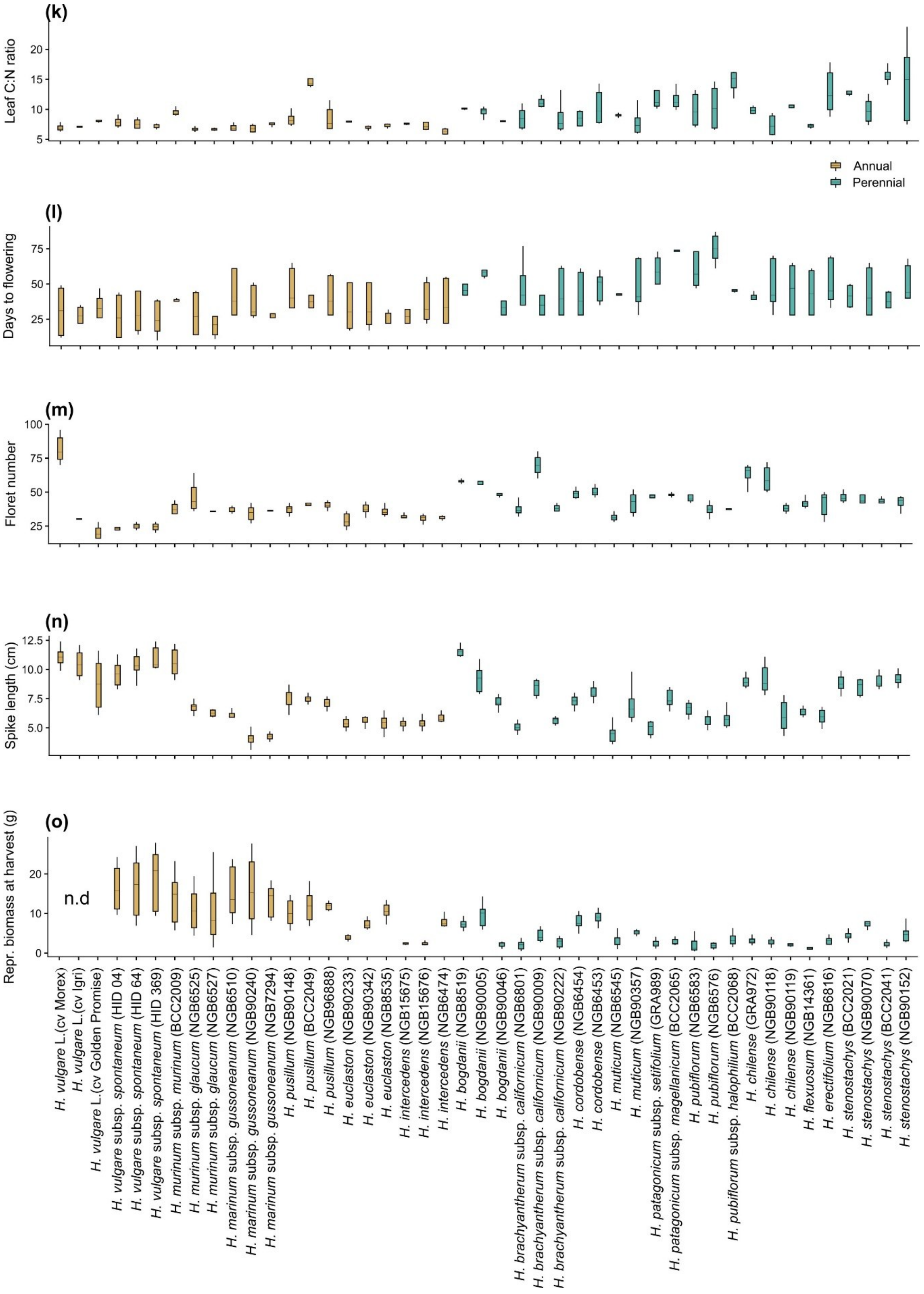
(k)
Leaf C:N ratio
Annual
Perennial
(l)
Days to flowering
(m)
Floret number
(n)
Spike length (cm)
(o)
Repr. biomass at harvest (g)
n.d
H. vulgare L.(cv Morex)
H. vulgare L.(cv Igri)
H. vulgare L.(cv Golden Promise)
H. vulgare subsp. spontaneum (HID 04)
H. vulgare subsp. spontaneum (HID 64)
H. vulgare subsp. spontaneum (HID 369)
H. murinum subsp. murinum (BCC2009)
H. murinum subsp. glaucum (NGB6525)
H. murinum subsp. glaucum (NGB6527)
H. marinum subsp. gussoneanum (NGB6510)
H. marinum subsp. gussoneanum (NGB90240)
H. marinum subsp. gussoneanum (NGB7294)
H. pusillum (NGB90148)
H. pusillum (BCC2049)
H. pusillum (NGB96888)
H. euclaston (NGB90233)
H. euclaston (NGB90342)
H. euclaston (NGB8535)
H. intercedens (NGB15675)
H. intercedens (NGB15676)
H. intercedens (NGB6474)
H. bogdanii (NGB8519)
H. bogdanii (NGB90005)
H. bogdanii (NGB90046)
H. brachyantherum subsp. californicum (NGB6801)
H. brachyantherum subsp. californicum (NGB90009)
H. brachyantherum subsp. californicum (NGB90222)
H. cordobense (NGB6454)
H. cordobense (NGB6453)
H. muticum (NGB6545)
H. muticum (NGB90357)
H. patagonicum subsp. setifolium (GRA989)
H. patagonicum subsp. magellanicum (BCC2065)
H. pubiflorum (NGB6583)
H. pubiflorum (NGB6576)
H. pubiflorum subsp. halophilium (BCC2068)
H. chilense (GRA972)
H. chilense (NGB90118)
H. chilense (NGB90119)
H. flexuosum (NGB14361)
H. erectifolium (NGB6816)
H. stenostachys (BCC2021)
H. stenostachys (NGB90070)
H. stenostachys (BCC2041)
H. stenostachys (NGB90152)

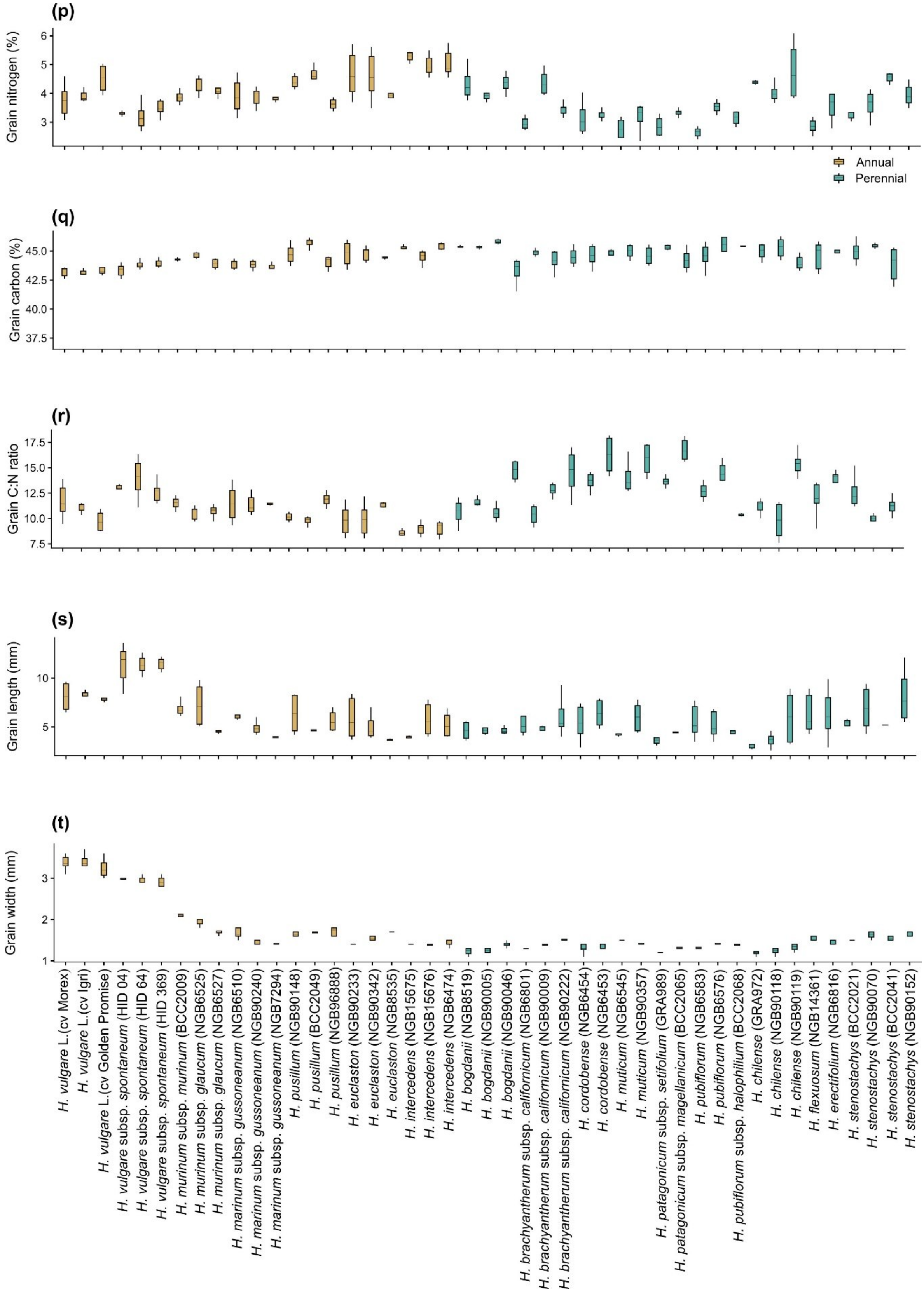
(p)
Grain nitrogen (%)
6
5
4
3
Annual
Perennial
(q)
Grain carbon (%)
45.0
42.5
40.0
37.5
(r)
Grain C:N ratio
17.5
15.0
12.5
10.0
7.5
(s)
Grain length (mm)
10
5
(t)
Grain width (mm)
3
2
1
H. vulgare L.(cv Morex)
H. vulgare L.(cv Igri)
H. vulgare L.(cv Golden Promise)
H. vulgare subsp. spontaneum (HID 04)
H. vulgare subsp. spontaneum (HID 64)
H. vulgare subsp. spontaneum (HID 369)
H. murinum subsp. murinum (BCC2009)
H. murinum subsp. glaucum (NGB6525)
H. murinum subsp. glaucum (NGB6527)
H. marinum subsp. gussoneanum (NGB6510)
H. marinum subsp. gussoneanum (NGB90240)
H. marinum subsp. gussoneanum (NGB7294)
H. pusillum (NGB90148)
H. pusillum (BCC2049)
H. pusillum (NGB96888)
H. euclaston (NGB90233)
H. euclaston (NGB90342)
H. euclaston (NGB8535)
H. intercedens (NGB15675)
H. intercedens (NGB15676)
H. intercedens (NGB6474)
H. bogdanii (NGB8519)
H. bogdanii (NGB90005)
H. bogdanii (NGB90046)
H. brachyantherum subsp. californicum (NGB6801)
H. brachyantherum subsp. californicum (NGB90009)
H. brachyantherum subsp. californicum (NGB90222)
H. cordobense (NGB6454)
H. cordobense (NGB6453)
H. muticum (NGB6545)
H. muticum (NGB90357)
H. patagonicum subsp. setifolium (GRA989)
H. patagonicum subsp. magellanicum (BCC2065)
H. pubiflorum (NGB6583)
H. pubiflorum (NGB6576)
H. pubiflorum subsp. halophilium (BCC2068)
H. chilense (GRA972)
H. chilense (NGB90118)
H. chilense (NGB90119)
H. flexuosum (NGB14361)
H. erectifolium (NGB6816)
H. stenostachys (BCC2021)
H. stenostachys (NGB90070)
H. stenostachys (BCC2041)
H. stenostachys (NGB90152)

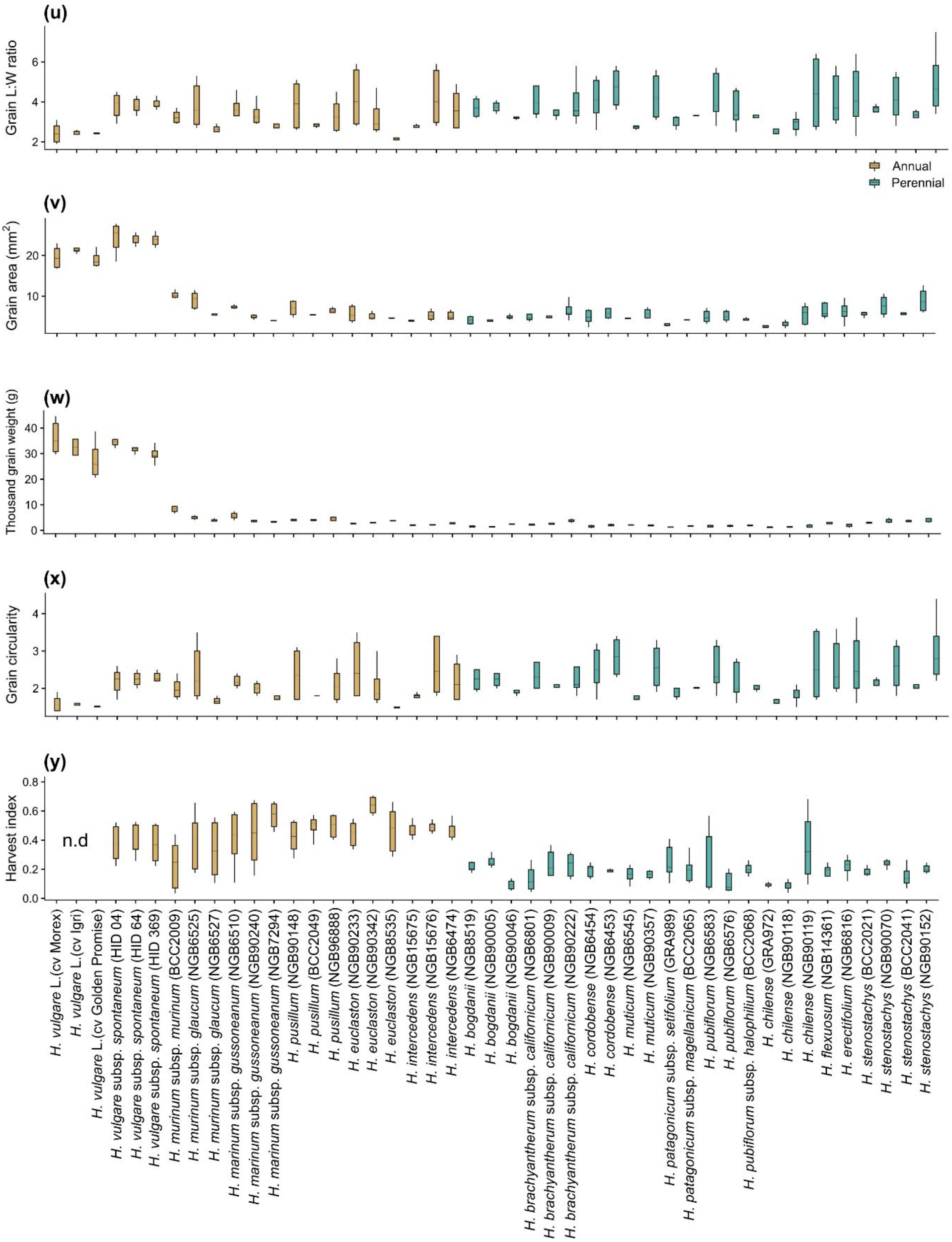


**Fig. S4: Boxplot showing mean trait values per accession, averaged across the 2021-2023 growing seasons (n = 8-12).**

Life history categories of accessions are indicated in brown (annuals, n = 21) and teal (perennials, n = 24). For each accession (x-axis), the genus *Hordeum is abbreviated as H* and the unique identifiers for each accession are in brackets. Refer to Table S1 for the full accession names. Dry shoot biom. 30-DAP, Dry shoot biomass after 30 days of planting (DAP); Leaf C: N, Flag leaf carbon-to-nitrogen ratio; Repr. biomass at harvest, Reproductive biomass at harvest; Grain C: N, Grain carbon-to-nitrogen ratio; Grain L: W, Grain length-to-width ratio. Leaf traits, including leaf area, length, width, specific leaf area, leaf C, and N, were quantified using the flag leaf. Accessions with n.d. indicate missing data.

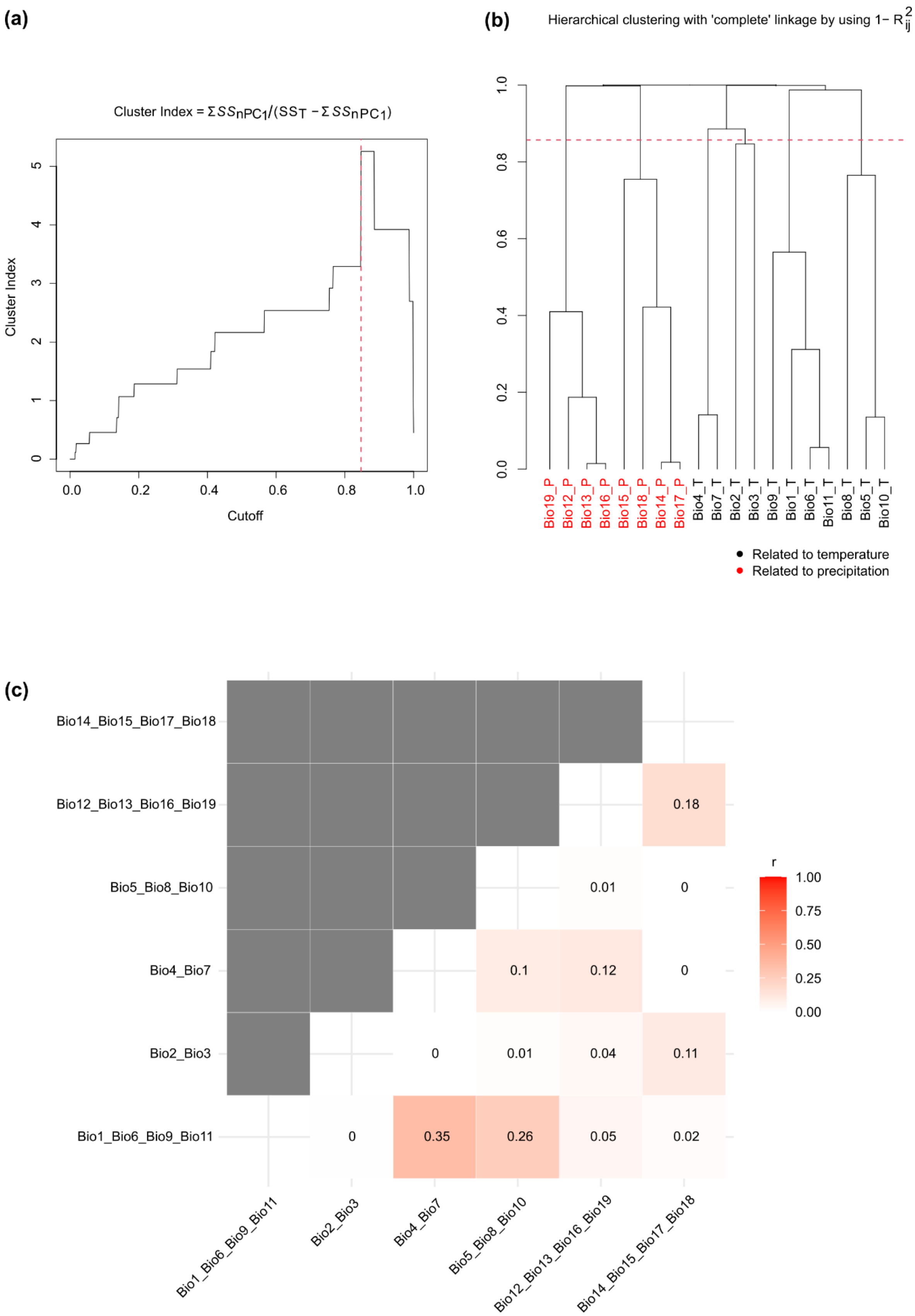


**Fig. S5: Procedure for generating synthetic environmental variables based on the methods described by Chang et al. (2022).**

**(a)** Relationship between clustering cutoff and corresponding cluster index. The y-axis represents the customized cluster index, and the x-axis represents $1 - R^2$, which serves as the cutoff for the dendrogram. The vertical red dashed line indicates the optimal cutoff. **(b)** Dendrogram of hierarchical clustering based on the $1 - R^2$. The horizontal red dashed line marks the optimal cutoff used to determine cluster combinations for generating synthetic environmental variables. **(c)** Correlation matrix of the six retained environmental variables. Abbreviations: Bio3_T, Isothermality; Bio4_T, Temperature seasonality; Bio8_T, Mean temperature of wettest quarter; Bio9_T, Mean temperature of driest quarter; Bio17_P, Precipitation of driest quarter; Bio19_P, Precipitation of coldest quarter.

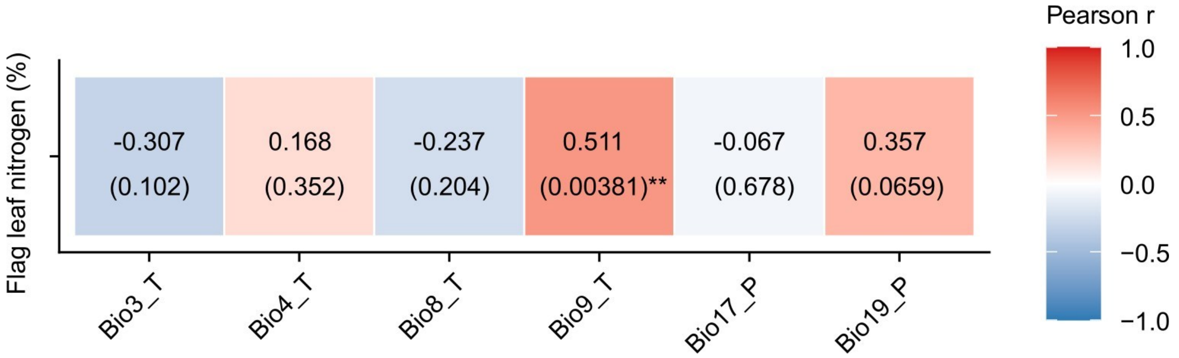


**Fig. S6: Associations between traits and climatic variables along the first significant constrained RDA axis after accounting for phylogenetic relatedness.**

Pairwise trait-climatic variables correlations (FDR-adjusted *p*-values) were calculated using Pearson correlation coefficients. The values in the heatmap represent trait-climatic variable correlations (outside brackets) and FDR-adjusted *p*-values (in brackets), respectively. Associations with FDR-adjusted $p$-values < 0.01 were considered significant. Abbreviations: Bio3_T, Isothermality; Bio4_T, Temperature seasonality; Bio8_T, Mean temperature of wettest quarter; Bio9_T, Mean temperature of driest quarter; Bio17_P, Precipitation of driest quarter; Bio19_P, Precipitation of coldest quarter. Significance levels: **, $p < 0.01$.

**Supplementary Tables**

**Table S1** List of *Hordeum* accessions used in the study. Columns include accession entry number (No.), accession names (AN), latitude (Lat), longitude (Lon), geographical origin (GO), genebank from which the seed originated (SO), accession identity (AI), life history (LH, A = Annual, P = Perennial), and years evaluated (YE). For perennial accessions, records of regrowth potential after winter are documented between experimental seasons. For those planted in 2020, the number of seasons survived (where verified) is shown in brackets next to LH.

| No. | AN | Lat | Lon | GO | SO | AI | LH | YE |
|---|---|---|---|---|---|---|---|---|
| 1 | *Hordeum marinum* subsp. *gussoneanum* (Parl.) Thell. | 32.90 | 40.10 | Ankara, Turkey | NordGen[1] | NGB6510 | A | 2021-2023 |
| 2 | *Hordeum marinum* subsp. *gussoneanum* (Parl.) Thell. | 39.3 | -5.5 | Lagrosan, Spain | BordGen[1] | NBG7294 | A | 2022, 2023 |
| 3 | *Hordeum marinum* subsp. *gussoneanum* (Parl.) Thell. | 38.4 | 23.8 | Athens, Greece | NordGen[1] | NGB90240 | A | 2021-2023 |
| 4 | *Hordeum murinum* subsp. *murinum* L. | 54.7 | 11.5 | Fyn, Denmark | IPK[2] | BCC2009 | A | 2022, 2023 |
| 5 | *Hordeum murinum* subsp. *glaucum* (Steud.) Tzvelev. | 38.4 | 27.1 | Izmir, Turkey | NordGen[1] | NGB6525 | A | 2021-2023 |
| 6 | *Hordeum murinum* subsp. *glaucum* (Steud.) Tzvelev. | 39 | 1.3 | Ibiza, Spain | NordGen[1] | NGB6527 | A | 2021-2023 |
| 7 | *Hordeum vulgare* subsp. *spontaneum* (K. Koch) Thell. | 35.5 | 43 | Mosul, Iraq | CREA[3] | HID 04 | A | 2021-2023 |
| 8 | *Hordeum vulgare* subsp. *spontaneum* (K. Koch) Thell. | 37.4 | 36.6 | Gaziantep, Turkey | CREA[3] | HID 64 | A | 2021-2023 |
| 9 | *Hordeum vulgare* subsp. *spontaneum* (K. Koch) Thell. | 32.9 | 35.4 | Fertile Crescent, Israel | CREA[3] | HID 369 | A | 2021-2023 |
| 10 | *Hordeum vulgare* L. | - | - | NA | NA | cv Morex | A | 2021-2023 |
| 11 | *Hordeum vulgare* L. | - | - | NA | NA | cv Golden Promise | A | 2022, 2023 |

| No. | AN | Lat | Lon | GO | SO | AI | LH | YE |
|---|---|---|---|---|---|---|---|---|
| 12 | *Hordeum vulgare* L. | - | - | NA | NA | cv Igri | A | 2022, 2023 |
| 13 | *Hordeum intercedens* Nevski | 34 | -117.4 | California, USA | NordGen[1] | NGB6474 | A | 2021-2023 |
| 14 | *Hordeum intercedens* Nevski | 30.4 | -115.9 | Baja California North, Mexico | NordGen[1] | NBG15675 | A | 2022, 2023 |
| 15 | *Hordeum intercedens* Nevski | 30.4 | -115.9 | Baja California North, Mexico | NordGen[1] | NGB15676 | A | 2021-2023 |
| 16 | *Hordeum pusillum* Nutt. | 39.7 | -105.3 | New Mexico, USA | IPK[2] | BCC2049 | A | 2022, 2023 |
| 17 | *Hordeum pusillum* Nutt. | 39.8 | -105.2 | Colorado, Denver, USA | NordGen[1] | NGB6888 | A | 2021-2023 |
| 18 | *Hordeum pusillum* Nutt. | 32.2 | -107.8 | New Mexico, USA | NordGen[1] | NGB90148 | A | 2021-2023 |
| 19 | *Hordeum euclaston* Steud. | -47.2 | -67.0 | Province Santa Cruz, Argentina | NordGen[1] | NGB8535 | A | 2022, 2023 |
| 20 | *Hordeum euclaston* Steud. | -32.2 | -57.9 | Paysandú, Uruguay | NordGen[1] | NGB90233 | A | 2021-2023 |
| **21** | ***Hordeum euclaston* Steud**. | **-47.1** | **-67.0** | **Santa Cruz, Argentina** | **NordGen[1]** | **NGB90342** | **A** | **2021-2023** |
| 22 | *Hordeum bogdanii* Wilensky | 33.0 | 69.2 | Paktia, Afghanistan | NordGen[1] | NGB8519 | P | 2022, 2023 |
| 23 | *Hordeum bogdanii* Wilensky | 39.5 | 75.9 | Xinjiang, China | NordGen[1] | NGB90005 | P (2) | 2022, 2023 |
| **24** | ***Hordeum bogdanii* Wilensky** | **35.8** | **52.7** | **Markazi, Iran** | **NordGen[1]** | **NGB90046** | **P (2)** | **2022, 2023** |
| 25 | *Hordeum brachyantherum* subsp. *californicum* (Covas & Stebbins) Bothmer, N. Jacobsen & Seberg | 37.4 | -120 | California, USA | NordGen[1] | NGB6801 | P (2) | 2021-2023 |
| 26 | *Hordeum brachyantherum* subsp. *californicum* (Covas & Stebbins) Bothmer, N. Jacobsen & Seberg | 36.8 | -119.8 | California, USA | NordGen[1] | NGB90009 | P | 2022, 2023 |

| No. | AN | Lat | Lon | GO | SO | AI | LH | YE |
|---|---|---|---|---|---|---|---|---|
| 27 | *Hordeum brachyantherum* subsp. *Californicum* (Covas & Stebbins) Bothmer, N. Jacobsen & Seberg | 32.7 | -117.2 | California, USA | NordGen[1] | NGB90222 | P | 2021-2023 |
| **28** | ***Hordeum chilense* Roem. & Schult.** | **-32.5** | **-71.4** | **Aconcagua, Chile** | **IPK[2]** | **GRA972** | **P** | **2022, 2023** |
| **29** | ***Hordeum chilense* Roem. & Schult.** | **-38.9** | **-70.1** | **Aconcagua, Chile** | **NordGen[1]** | **NGB90118** | **P (2)** | **2021-2023** |
| 30 | *Hordeum chilense* Roem. & Schult. | -30.0 | -71.3 | Coquimbo, Chile | NordGen[1] | NGB90119 | P (2) | 2021-2023 |
| 31 | *Hordeum cordobense* Bothmer, N. Jacobsen & Nicora | -31.3 | -64.2 | Cordoba, Argentina | NordGen[1] | NGB6453 | P (3) | 2021-2023 |
| 32 | *Hordeum cordobense* Bothmer, N. Jacobsen & Nicora | -31.3 | -64.2 | Cordoba, Argentina | NordGen[1] | NGB6454 | P (3) | 2021-2023 |
| 33 | *Hordeum erectifolium* Bothmer, N. Jacobsen & R.B. Jørg. | -38.5 | -62.3 | Buenos Aires, Argentina | NordGen[1] | NGB6816 | P (3) | 2021-2023 |
| 34 | *Hordeum flexuosum* Nees ex Steud. | -34.4 | -58.5 | Buenos Aires, Argentina | NordGen[1] | NGB14361 | P (2) | 2021-2023 |
| 35 | *Hordeum muticum* J. Presl | -17.4 | -66.1 | Cochabamba, Bolivia | NordGen[1] | NGB6545 | P (2) | 2022, 2023 |
| 36 | *Hordeum muticum* J. Presl | -24.9 | -65.4 | Salta, Argentina | NordGen[1] | NGB90357 | P (2) | 2021-2023 |
| 37 | *Hordeum stenostachys* Godr. | -34.6 | -58.5 | Buenos Aires, Argentina | IPK[2] | BCC2021 | P | 2022, 2023 |
| 38 | *Hordeum stenostachys* Godr. | -28.9 | -67.5 | La Rioja, Argentina | IPK[2] | BCC2041 | P | 2022, 2023 |
| 39 | *Hordeum stenostachys* Godr. | -29.4 | -68.7 | La Rioja, Argentina | NordGen[1] | NGB90152 | P (2) | 2021-2023 |
| 40 | *Hordeum stenostachys* Godr. | -31.5 | -68.5 | San Juan, Argentina | NordGen[1] | NGB90070 | P (2) | 2021-2023 |
| 41 | *Hordeum patagonicum* subsp. *setifolium* (Parodi ex Nicora) Bothmer, Giles & N. Jacobsen | -44.7 | -70.2 | Chubut, Argentina | IPK[2] | GRA989 | P (2) | 2022, 2023 |

| No. | AN | Lat | Lon | GO | SO | AI | LH | YE |
|---|---|---|---|---|---|---|---|---|
| 42 | *Hordeum patagonicum* subsp. *magellanicum* (Parodi ex Nicora) Bothmer, Giles & N. Jacobsen | -51.6 | -69.2 | Santa Cruz, Argentina | IPK[2] | BCC2065 | P (2) | 2022, 2023 |
| 43 | *Hordeum pubiflorum* subsp. *halophilum* (Griseb.) C. Baden & Bothmer | -19.5 | -65.7 | Quijarro, Bolivia | IPK[2] | BCC2068 | P | 2022, 2023 |
| 44 | *Hordeum pubiflorum* Hook. f. | -43.4 | -70.8 | Chubut, Argentina | NordGen[1] | NGB6583 | P (2) | 2021-2023 |
| 45 | *Hordeum pubiflorum* Hook. f. | -50.4 | -72.5 | Santa Cruz, Argentina | NordGen[1] | NGB6576 | P (2) | 2021-2023 |

[1]Nordic Genetic Resource Center, Sweden

[2]Leibnitz-Institut für Pflanzengenetik und Kulturpflanzenforschung, Gatersleben, Germany

[3]Centro di Ricerca Genomica e Bioinformatica, sede di Fiorenzuola d′Arda, Italy

Accessions with entry numbers 21, 24, 28, and 29 were excluded from the RDA because they were not included in the accession-based phylogenetic tree published by Hellwig et al. (2025).

NA: geo-referenced coordinates could not be generated and were thus also excluded from the RDA.

cv: cultivated variety.

**Table S2** Air temperature (ºC) with mean standard deviation (±SD) for day, night, and overall, by month and year. The values are daily recordings averaged monthly for the five-month-long common garden experiment.

| Year | Month | Day | Night | Mean |
|---|---|---|---|---|
| 2021 | April | 10.8 ± 4.7 | 7.2 ± 4.3 | 9.0 ± 4.8 |
| | May | 14.0 ± 4.6 | 11.0 ± 4.2 | 12.5 ± 4.6 |
| | June | 22.6 ± 5.1 | 18.7 ± 4.5 | 20.7 ± 5.2 |
| | July | 21.0 ± 3.7 | 17.6 ± 3.0 | 19.3 ± 3.7 |
| | August | 19.4 ± 3.8 | 16.3 ± 2.9 | 17.9 ± 3.7 |
| 2022 | April | 14.9 ± 4.5 | 10.7 ± 3.6 | 12.8 ± 4.6 |
| | May | 18.9 ± 5.1 | 14.3 ± 4.4 | 16.6 ± 5.3 |
| | June | 21.5 ± 4.9 | 17.3 ± 4.7 | 19.4 ± 5.3 |
| | July | 22.8 ± 5.1 | 18.8 ± 4.2 | 20.8 ± 5.1 |
| | August | 24.8 ± 5.3 | 20.4 ± 4.2 | 22.6 ± 5.3 |
| 2023 | April | 12.2 ± 4.0 | 9.1 ± 3.3 | 10.7 ± 4.0 |
| | May | 15.9 ± 3.9 | 12.1 ± 3.7 | 14.0 ± 4.3 |

Note: Daytime was defined as 05:00–21:00 h and nighttime as 21:00–5:00 h, based on local German time. Data for air temperature between June and August for the 2023 season could not be generated due to a faulty watchdog.

**Table S3** Description of phenotypic traits, functional importance, references, and the years in which traits were evaluated.

| Trait group | Trait | Description | Functional importance | Reference | Year |
|---|---|---|---|---|---|
| Vegetative | Flag leaf area ($cm^2$) | Measured using the *Petiole* mobile application (non-pro version, first downloaded on 05.05.2021) | Proxy for heat load, water retention, and gas exchange | (Stubbs & Wilson, 2004; Yang et al., 2019) | 2021-2023 |
| | Flag leaf length (cm) | Manually measured with a folding rule from the leaf collar to the tip of the blade | Gas exchange, heat load | (Stubbs & Wilson, 2004) | 2021-2023 |
| | Flag leaf width (cm) | Manually measured with a folding rule at the widest section of the leaf blade | Gas exchange, heat load | (Stubbs & Wilson, 2004) | 2021-2023 |
| | Plant height (cm) | Manually measured from the base of the main tiller (at the soil level) to the collar of the spike | Reflects plant productivity, nutrient availability, competition for light, and growth strategy | (Moles et al., 2009; Li & Prentice, 2024) | 2021-2023 |
| | Dry shoot biomass after 30 days (g) | Whole shoot biomass aboveground was cut and dried in paper bags for 72 hrs in a fan-assisted oven set to 65 ºC before weighing | Proxy for plant productivity (growth rate) | (Quétier et al., 2007; Zieschank & Junker, 2023) | 2022-2023 |
| | Stem diameter at the base (mm) | Measured at harvest on the mid-sections between the first and second nodes | Reflects growth potential and mortality risk from biomechanical or hydraulic failure | (Díaz et al., 2016) | 2021, 2023 |

| Trait group | Trait | Description | Functional importance | Reference | Year |
|---|---|---|---|---|---|
| | | above the soil surface using a digital caliper | | | |
| | Shoot biomass at harvest (g) | Dry biomass of the whole shoot aboveground was weighed post-harvest | Proxy for plant productivity | (Quétier et al., 2007; Zieschank & Junker, 2023) | 2022-2023 |
| | Specific leaf area ($cm^2$ $mg^{-1}$) | Specific leaf area = $\frac{flag\ leaf\ area\ (cm2)}{\text{weight of dry leaf area (mg)}}$ | Reflects photosynthetic capacity, leaf longevity, and stress tolerance | (Westoby, 1999) | 2021-2023 |
| | Flag leaf nitrogen (%) | Measurement was performed on homogenized flag leaf powder using a mass spectrometer coupled to an isotope cube elemental analyzer | Reflect photosynthetic potential, leaf longevity | (Reich et al., 1991) | 2021-2022 |
| | Flag leaf carbon (%) | Measurement was performed on homogenized flag leaf powder using a mass spectrometer coupled to an isotope cube elemental analyzer | Reflects energy transfer, plant growth, and productivity | (Bruelheide et al., 2018; Li & Prentice, 2024) | 2021-2022 |
| | Flag leaf carbon-to-nitrogen ratio | Value or ratio generated from carbon and nitrogen measurements | Energy transfer, plant growth, productivity, and photosynthetic potential | (Bruelheide et al., 2018; Li & Prentice, 2024) | 2021-2022 |

| Trait group | Trait | Description | Functional importance | Reference | Year |
|---|---|---|---|---|---|
| Reproductive | Days to flowering | Determined as the number of days from repotting to the outdoors until the first awns from the flag leaf appeared | Proxy for fitness, yield | (Gaudinier & Blackman, 2020) | 2021-2023 |
| | Floret number | Manually counted using a digital hand tally counter | Proxy for yield | (Alqudah & Schnurbusch, 2014; Alqudah et al., 2021; Liu et al., 2022) | 2021-2022 |
| | Spike length (cm) | Manually measured from the collar of the spike to the tip, excluding the awns that extend beyond the spike | Proxy for yield | (Alqudah & Schnurbusch, 2014; Alqudah et al., 2021; Liu et al., 2022) | 2021-2023 |
| | Reproductive biomass (g) | Represents the weight of spikes measured after harvest | Measures reproductive efficiency | (Unkovich et al., 2010) | 2022-2023 |
| | Grain nitrogen (%) | Measurement was performed on homogenized grain powder using a mass spectrometer coupled to an isotope cube elemental analyzer | Reflects nitrogen remobilization from vegetative tissues, energy transfer, and productivity | (Gaju et al., 2014; Kong et al., 2016) | 2021-2022 |

| Trait group | Trait | Description | Functional importance | Reference | Year |
|---|---|---|---|---|---|
| | Grain carbon (%) | Measurement was performed on homogenized grain powder using a mass spectrometer coupled to an isotope cube elemental analyzer | Reflects energy transfer and productivity | (Gaju et al., 2014; Kong et al., 2016) | 2021-2022 |
| | Grain carbon-to-nitrogen ratio | Value or ratio generated from carbon and nitrogen measurements | Reflects energy transfer and productivity | (Gaju et al., 2014; Kong et al., 2016) | 2021-2022 |
| | Grain length (mm) | Determined using the MARVIN Seed Analyzer from 30-50 grains, finely cleaned | Reflects dispersal, regeneration, and reproduction ability | (Bruelheide et al., 2018) | 2021-2023 |
| | Grain width (mm) | Determined using the MARVIN Seed Analyzer from 30-50 grains, finely cleaned | Reflects dispersal, regeneration, and reproduction ability | (Bruelheide et al., 2018) | 2021-2023 |
| | Grain length-to-width ratio | Determined using the MARVIN Seed Analyzer from 30-50 grains, finely cleaned | Reflects dispersal, regeneration, and reproduction ability | (Bruelheide et al., 2018) | 2021-2023 |
| | Grain area ($mm^2$) | Determined using the MARVIN Seed Analyzer from 30-50 grains, finely cleaned | Reflects dispersal, regeneration, and reproduction ability | (Bruelheide et al., 2018) | 2021-2023 |
| | Thousand-grain weight (g) | Determined using the MARVIN Seed Analyzer | Reflects dispersal, regeneration, and reproduction ability | (Moles & Westoby, | 2021-2023 |

| Trait group | Trait | Description | Functional importance | Reference | Year |
|---|---|---|---|---|---|
| | | from 30-50 grains, finely cleaned | | 2006; Díaz et al., 2016) | |
| | Grain circularity | Determined using the MARVIN Seed Analyzer from 30-50 grains, finely cleaned | Reflects dispersal, regeneration, and reproduction ability | (Bruelheide et al., 2018) | 2021-2023 |
| | Harvest index | Harvest index = $\frac{Spike\ biomass\ at\ harvest\ (g)}{\text{dry shoot biomass at harvest (g)}}$ | Measure of reproductive efficiency | (Unkovich et al., 2010) | 2022-2023 |

**Table S5** All environmental variables retrieved from the WorldClim2 database and used in spatial niche modelling to identify the variables that best describe the niches of the different *Hordeum* accessions.

| Abbreviation | Description | Unit |
|---|---|---|
| Bio1_T | Annual mean temperature | ºC |
| Bio2_T | Mean diurnal range (mean of monthly (max temp – min temp)) | ºC |
| Bio3_T | **Isothermality (bio2/bio7) (x 100)** | % |
| Bio4_T | **Temperature seasonality (standard deviation x 100)** | % |
| Bio5_T | Maximum temperature of warmest month | ºC |
| Bio6_T | Minimum temperature of coldest month | ºC |
| Bio7_T | Temperature annual range (Bio5-bio6) | ºC |
| Bio8_T | **Mean temperature of wettest quarter** | ºC |
| Bio9_T | **Mean temperature of driest quarter** | ºC |
| Bio10_T | Mean temperature of warmest quarter | ºC |
| Bio11_T | Mean temperature of coldest quarter | ºC |
| Bio12_P | Annual precipitation | mm |
| Bio13_P | Precipitation of the wettest month | mm |
| Bio14_P | Precipitation of the driest month | mm |
| Bio15_P | Precipitation seasonality (coefficient of variation) | % |
| Bio16_P | Precipitation of the wettest quarter | mm |
| Bio17_P | **Precipitation of the driest quarter** | mm |
| Bio18_P | Precipitation of the warmest quarter | mm |
| Bio19_P | **Precipitation of the coldest quarter** | mm |

Environmental variables selected for trait partitioning in the RDA models are in bold.

**Table S6** Comparison of traits between annual and perennial species after accounting for phylogeny. Traits are ranked in descending order of partial $R^2$. Trait values are presented as mean ± standard deviation (SD). Effect sizes are reported as Cohen's *d* with 95% confidence intervals; positive values indicate higher mean values in annuals and vice versa.

| Trait | Mean (Annual ± SD) | Mean (Perennial ± SD) | Effect size (Cohen's *d* [95% CI]) | Effect magnitude | Significance | $R^2$ (%) |
|---|---|---|---|---|---|---|
| Harvest index | 0.07 ± 0.08 | -0.06 ± 0.04 | 1.95 [1.30, 3.25] | Large | *** | 51.4 |
| Grain nitrogen (%) | 0.25 ± 0.41 | -0.24 ± 0.44 | 1.13 [0.53, 1.91] | Large | *** | 25.9 |
| Grain C: N ratio | -0.81 ± 1.19 | 0.77 ± 1.58 | -1.12 [-1.83, -0.58] | Large | *** | 25.2 |
| Flag leaf carbon (%) | -0.14 ± 0.23 | 0.13 ± 0.25 | -1.07 [-1.81, -0.49] | Large | ** | 23.8 |
| Days to flowering (days) | -4.45 ± 6.62 | 4.24 ± 9.95 | -1.01 [-1.60, -0.47] | Large | *** | 21.5 |
| Grain L: W ratio | -0.08 ± 0.21 | 0.08 ± 0.15 | -0.85 [-1.45, -0.29] | Large | ** | 16.5 |
| Grain circularity | -0.05 ± 0.13 | 0.05 ± 0.10 | -0.84 [-1.51, -0.34] | Large | ** | 16.2 |
| Flag leaf C: N ratio | -0.40 ± 0.91 | 0.38 ± 0.98 | -0.80 [-1.49, -0.22] | Large | ** | 14.9 |
| Flag leaf nitrogen (%) | 0.20 ± 0.51 | -0.19 ± 0.47 | 0.79 [0.18, 1.50] | Medium | * | 14.7 |
| Grain width (mm) | 0.07 ± 0.18 | -0.06 ± 0.15 | 0.78 [0.22, 1.50] | Medium | * | 14.3 |

| Trait | Mean (Annual ± SD) | Mean (Perennial ± SD) | Effect size (Cohen's *d* [95% CI]) | Effect magnitude | Significance | $R^2$ (%) |
|---|---|---|---|---|---|---|
| Reproductive biomass at harvest (g) | 1.05 ± 3.38 | -1.00 ± 2.17 | 0.71 [0.12, 1.53] | Medium | * | 12.3 |
| Plant height (cm) | -5.04 ± 9.43 | 4.80 ± 17.28 | -0.69 [-1.33, -0.14] | Medium | * | 11.5 |
| Thousand grain weight (g) | 0.90 ± 3.09 | -0.86 ± 1.85 | 0.68 [0.07, 1.56] | Medium | * | 11.2 |
| Floret number | -3.00 ± 11.39 | 2.86 ± 6.68 | -0.62 [-2.02, 0.07] | Medium | ns | 9.5 |
| Stem diameter at base (mm) | -0.17 ± 0.52 | 0.16 ± 0.54 | -0.62 [-1.34, -0.04] | Medium | * | 9.4 |
| Dry shoot biomass (g) | 0.71 ± 3.22 | -0.67 ± 1.56 | 0.54 [-0.05, 1.19] | Medium | ns | 7.4 |
| Spike length (cm) | -0.36 ± 1.15 | 0.34 ± 1.71 | -0.47 [-1.11, 0.17] | Small | ns | 5.6 |
| Grain area ($mm^2$) | 0.23 ± 1.61 | -0.22 ± 1.38 | 0.29 [-0.31, 1.08] | Small | ns | 2.3 |
| Flag leaf width (cm) | 0.02 ± 0.17 | -0.02 ± 0.13 | 0.27 [-0.36, 0.91] | Small | ns | 2.0 |
| Shoot biomass at harvest (g) | -1.33 ± 7.67 | 1.27 ± 11.27 | -0.26 [-0.86, 0.39] | Small | ns | 1.9 |
| Grain length (mm) | -0.08 ± 0.83 | 0.07 ± 0.63 | -0.20 [-0.80, 0.44] | Negligible | ns | 1.1 |

| Trait | Mean (Annual ± SD) | Mean (Perennial ± SD) | Effect size (Cohen's *d* [95% CI]) | Effect magnitude | Significance | $R^2$ (%) |
|---|---|---|---|---|---|---|
| Flag leaf length (cm) | -0.19 ± 1.72 | 0.18 ± 2.78 | -0.16 [-0.76, 0.50] | Negligible | ns | 0.7 |
| Specific leaf area ($cm^2\ g^{-1}$) | -0.00 ± 0.00 | 0.00 ± 0.00 | -0.10 [-0.72, 0.55] | Negligible | ns | 0.3 |
| Flag leaf area ($cm^2$) | 0.07 ± 2.78 | -0.06 ± 2.10 | 0.05 [-0.66, 0.62] | Negligible | ns | 0.1 |
| Grain carbon (%) | 0.00 ± 0.24 | -0.00 ± 0.22 | 0.01 [-0.60, 0.69] | Negligible | ns | 0.0 |

Significance levels: ns, $p > 0.1$; *, $p < 0.05$; **, $p < 0.01$; ***, $p < 0.001$.

**Table S7** Trait-environment associations.

| Model | Type | Variance (%) | *p*-value |
|---|---|---|---|
| Trait ~ Env | Simple | 33 | 0.001*** |
| Trait ~ Env \| (Phy) | Partial | 15 | 0.012* |

Significance tested with 5000 permutations (ANOVA). Significance levels: *, $p < 0.05$; ***, $p < 0.001$.

**Table S8** Effects of individual climatic variables estimated by RDA models.

| Model type | Environmental variable | *F*-value | Var | Var (%) | *p*-value |
|---|---|---|---|---|---|
| [c]Marginal effect of climatic variables conditioned on phylogeny | Bio9_T | 3.34 | 1.40 | 5.61 | 0.0088** |
| | Bio8_T | 2.74 | 1.15 | 4.60 | 0.0208* |
| | Bio3_T | 1.42 | 0.60 | 2.38 | 0.1962ns |
| | Bio17_P | 1.41 | 0.59 | 2.37 | 0.2050ns |
| | Bio4_T | 1.28 | 0.54 | 2.16 | 0.2470ns |
| | Bio19_P | 1.07 | 0.45 | 1.80 | 0.3427ns |

Individual climatic variables are arranged according to explained trait variation (highest to lowest). Permutation tests computed *p*-values based on 5,000 permutations. Significance levels: ns, $p > 0.1$; *, $p < 0.05$; **, $p < 0.01$. Abbreviations of environmental variables are as follows: Bio3_T: Isothermality (Bio2/Bio7) (x 100), Bio4_T: Temperature seasonality (standard deviation x 100), Bio8_T: Mean temperature of wettest quarter, Bio9_T: Mean temperature of driest quarter, Bio17_P: Precipitation of driest quarter, Bio19_P: Precipitation of coldest quarter. Var: Variance

**Table S9** Trait-environment associations for all RDA axes. Analysis of variance (ANOVA) of each of the six constrained axes for significance.

| Model type | Axis | Variance | *F*-value | *p*-value |
|---|---|---|---|---|
| Partial RDA conditioned on phylogeny | RDA1 | 2.36 | 6.44 | 0.018* |
| | RDA2 | 0.65 | 1.79 | 0.76ns |
| | RDA3 | 0.32 | 0.88 | 0.99ns |
| | RDA4 | 0.27 | 0.73 | 0.98ns |
| | RDA5 | 0.11 | 0.31 | 0.99ns |
| | RDA6 | 0.09 | 0.24 | 0.99ns |
| | Residual | 11.35 | | |

Significance tested with 5000 permutations (ANOVA). Significance levels: ns, $p > 0.1$; *, $p < 0.05$

**Table S10** Candidate trait-environment associations detected only on the first significant constrained RDA axis after accounting for phylogenetic relatedness.

| Trait | Environmental variables | Correlation |
|---|---|---|
| Flag leaf area ($cm^2$) | Bio4_T | ns |
| Flag leaf length (cm) | Bio4_T | ns |
| Flag leaf width (cm) | Bio4_T | ns |
| Plant height (cm) | Bio8_T | ns |
| Dry shoot biomass after 30 days (g) | Bio19_P | ns |
| Stem diameter at the base at harvest (mm) | Bio4_T | ns |
| Shoot biomass at harvest (g) | Bio3_T | ns |
| Specific leaf area ($cm^2mg^{-1}$) | Bio3_T | ns |
| **Flag leaf nitrogen (%)** | **Bio9_T** | **0.51**** |
| Flag leaf carbon (%) | Bio9_T | ns |
| Flag leaf carbon-to-nitrogen ratio | Bio9_T | ns |
| Days to flowering | Bio9_T | ns |
| Floret number | Bio9_T | ns |
| Spike length (cm) | Bio4_T | ns |
| Reproductive biomass after harvest (g) | Bio4_T | ns |
| Grain nitrogen (%) | Bio17_P | ns |
| Grain carbon (%) | Bio3_T | ns |
| Grain carbon-to-nitrogen ratio | Bio9_T | ns |
| Grain length (mm) | Bio19_P | ns |
| Grain width (mm) | Bio19_P | ns |
| Grain length-to-width ratio | Bio8_T | ns |
| Grain area ($mm^2$) | Bio19_P | ns |
| Thousand-grain weight (g) | Bio19_P | ns |
| Grain circularity | Bio8_T | ns |
| Harvest index | Bio8_T | ns |

The candidate trait-environment association coefficient detected on the first significant constrained RDA axis is in bold. Abbreviations: Bio3_T, Isothermality (Bio2/Bio7) (x 100); Bio4_T, Temperature seasonality (standard deviation x 100); Bio8_T, Mean temperature of wettest quarter; Bio9_T, Mean temperature of driest quarter; Bio17_P, Precipitation of driest quarter; Bio19_P, Precipitation of coldest quarter. Significance levels: ns, $p > 0.1$; ***, $p < 0.001$.
